# Missense variants in human forkhead transcription factors reveal determinants of forkhead DNA bispecificity

**DOI:** 10.1101/2025.05.27.656303

**Authors:** Jessica King, Stephen S. Gisselbrecht, Julie-Alexia Dias, Raehoon Jeong, Elisabeth Rothman, Martha L. Bulyk

## Abstract

Recognition of specific DNA sequences by transcription factors (TFs) is a key step in transcriptional control of gene expression. While most forkhead (FH) TFs bind either an FKH (RYAAAYA) or an FHL (GACGC) recognition motif, some FHs can bind both motifs. Mechanisms that control whether a FH is monospecific versus bispecific have remained unknown. Screening a library of 12 reference FH proteins, 61 naturally occurring missense variants including clinical variants, and 22 designed mutant FHs for DNA binding activity using universal (“all 10-mer”) protein binding microarrays (PBMs) revealed non-DNA-contacting residues that control mono- versus bispecificity. Variation in non-DNA-contacting amino acid residues of TFs is associated with human traits and may play a role in the evolution of TF DNA binding activities and gene regulatory networks.

**Highlights:**

- Most forkhead (FH) proteins recognize FKH or FHL motifs, while others are bispecific
- DNA binding activities of 12 reference and 83 variant or designed FHs
- Clinical or population FH variants with altered DNA binding affinity or specificity
- Non-DNA-contacting amino acid residues synergistically control mono- versus bi- specificity

## Introduction

Binding by sequence-specific transcription factors (TFs) to their recognition sites is a primary step in gene regulation. TF mutations, especially those within TF DNA binding domains (DBDs), can alter TF DNA binding activity and cause differences in traits, both in disease ^1^ and evolution ^2–4^. Furthermore, different mutations within a particular TF DBD can have a spectrum of effects on DNA binding activity, including changes in the strength of DNA binding (“affinity”) or sequence specificity. Surveys of the effects of TF mutations on DNA binding activity have revealed TF DNA binding specificity determinants ^5–9^.

The Forkhead (FH) family of TFs is a functionally diverse group of sequence-specific DNA-binding factors ^10^, with multiple members in all animal and fungal species. Various human FH family members have crucial roles in the development of a wide range of cell types and organ systems, in cellular response to metabolic signals, and in control of proliferation. Inherited and/or somatic variation in forkhead TF genes has been associated with many different disease states ^11^; for example, various missense substitutions in FOXC1 cause developmental disorders, including ocular defects in Axenfeld-Rieger syndrome, through a range of pathogenic effects on transcriptional gene regulation, including altered DNA binding activity ^12^.

The FH family is defined by the presence of a FH DBD. This is a structurally compact domain belonging to the winged helix superfamily ^10^(Figure 1A). Winged helix domains consist of a compact 3-helix bundle with a hydrophobic core, similar to the homeodomain and other evolutionarily ancient helix-turn-helix DBDs, along with two “wings” ^13^. Wing 1 is a small (2- or 3- strand) beta sheet with a loop between the strands, while wing 2 is much less well conserved and adopts very different structures between family members. Some other families of winged helix proteins use the wings to make DNA specificity-determining contacts ^13^, but FH domains insert a recognition helix, helix 3, into the DNA major groove and use the wings to make stabilizing backbone contacts in all protein-DNA cocrystal structures examined to date ^14,15^. Highly conserved residues in the recognition helix determine DNA-binding specificity by making direct and water-mediated base-specific contacts.

**Figure 1:**
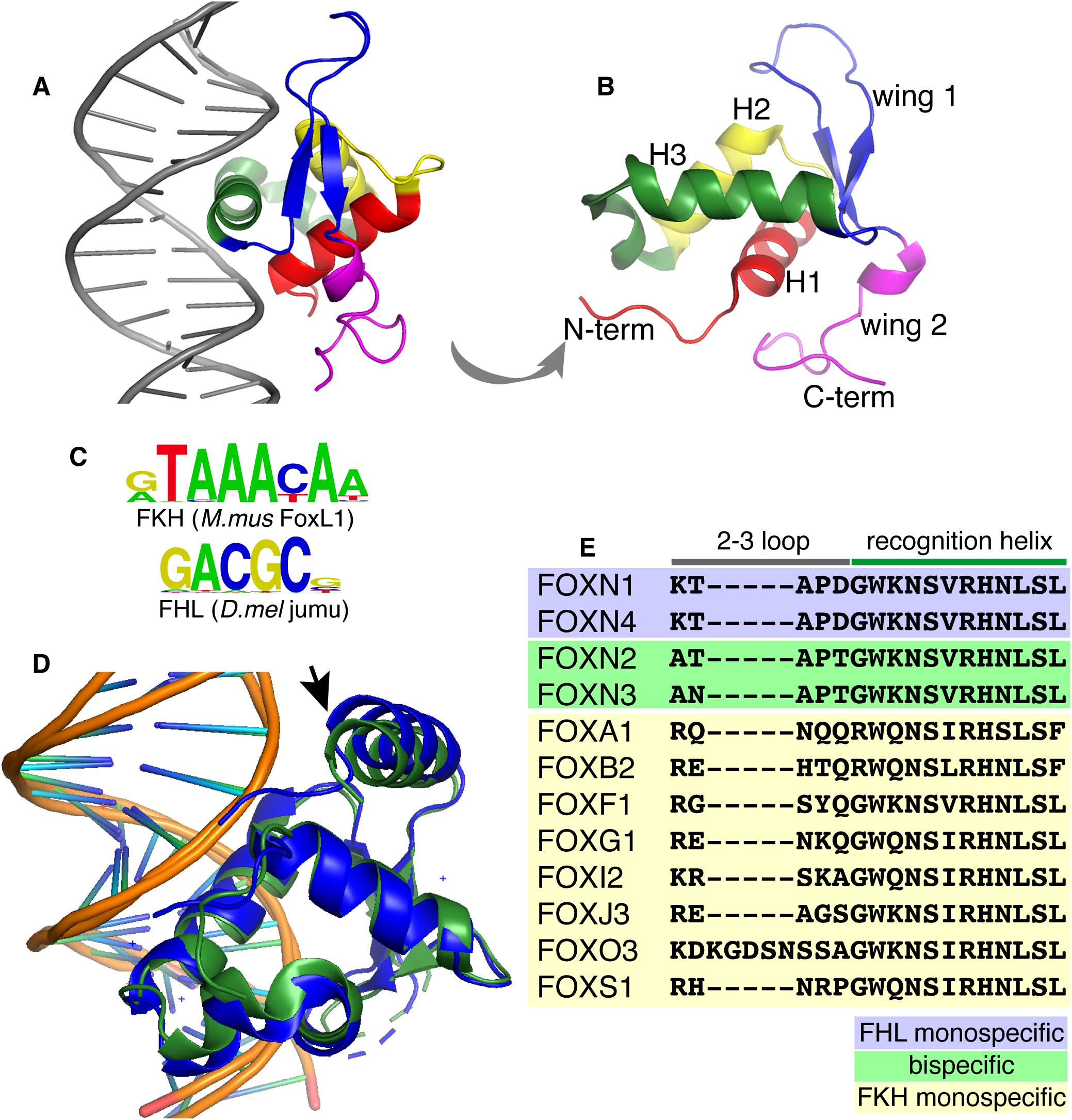
Current knowledge about the structure and specificity determinants of the FH domain. **(A)** Structure of a representative FH domain (FOXC2, PDB accession # 6AKO) bound to DNA, and **(B)**, as viewed from the DNA (rotated approximately 90 degrees). The N-terminal tail and helix 1 are shown in red, helix 2 in yellow, helix 3 and the 2-3 loop in green, wing 1 in blue, and wing 2 in purple. **(C)** Sequence logo representations of the FKH and FHL motif (from Nakagawa *et al.* ^20^). **(D)** Alignment of the structures of FOXN3 in complex with the FKH and FHL motifs, from Rogers *et al.* ^24^. The arrow indicates the portion of wing 2 that shows the largest divergence in protein structure between the two complexes. **(E)** Aligned protein sequences of the 2-3 loop and recognition helix for all RefSeq FH domains tested in this study, grouped by specificity.

The FH family in metazoans is divided into approximately 20 subfamilies, on the basis of shared protein sequence motifs within the domain ^16^. Despite this diversity within the family, it was long believed that all FH domains bind most specifically to a core RYAAAYA recognition motif and minor variants thereof, with different family members showing relatively subtle base preferences at the partially degenerate positions and flanking the core ^17^. Work in our and other labs has shown that many FH domains recognize this RYAAAYA motif, which we refer to as the canonical forkhead (“FKH”) motif, but that some subfamilies strongly prefer a very different, GACGC motif, which we term the “FHL” motif, corresponding to the recognition motif of the yeast *Saccharomyces cerevisiae* forkhead-like (Fhl1) protein (Figure 1B) ^18–20^. All eukaryotic TFs bind multiple sequences, but these are typically more similar to each other than are the FKH and FHL motifs, with minor variation at positions of lower information content within the recognition sequence. Large changes in specificity within a family are generally associated with variation in the amino acid residues that make base-selective contacts to mediate sequence specificity; however, we found that FH binding to FKH vs. FHL sequences does not correlate with these changes in the evolution of the FH family ^20^. Strikingly, we found that there are some “bispecific” FH domains that bind to both of these DNA sequence motifs with little apparent preference between them ^20^. Within the FOXN subfamily, FOXN2 and FOXN3 are bispecific, while FOXN1 and FOXN4 strongly prefer the FHL motif. FOXM1 (the only member of the FOXM subfamily) is bispecific for FKH binding and for a variant of the FHL motif (CATGC), and this alternate binding specificity appears to have arisen independently during the evolution of the family, suggesting that there is some inherent flexibility in the mechanism of FH-DNA recognition. Unsupervised clustering of experimentally determined *in vitro* DNA binding profiles partitioned the different FH domains into three broad specificity classes: FKH-monospecific, FHL-monospecific, and bispecific ^20^.

The discovery of bispecificity raised the question of what mediates recognition of two distinct classes of DNA binding sequences with radically different G/C content and different lengths. The availability of DNA-protein cocrystal structures for several FKH-monospecific FH domains (including FOXK1, FOXC2 and FOXG1 ^21–23^) and the FHL-monospecific FOXN1 FH domain, and for the bispecific FOXN3 domain bound separately to each of its cognate motifs, provided a partial answer. The same DBD contact surface and amino acid residues make different base-specific contacts when bound to the two different motif classes, with significant variation only in the side chain rotamer of a single ultra-conserved, base-contacting asparagine residue in the recognition helix. Instead, the principle difference observed is in the shape of the bound DNA molecule, which is bent away from the protein in FHL-bound structures but bends toward the protein to accommodate the longer, FKH motif. This discovery raised the question of why every FH domain does not bind to both motifs.

Two regions of the FH domain have received the most attention to date as candidate determinants of motif specificity. Rogers *et al.* used a subdomain-swap strategy to address this question: portions of the bispecific FH domain of FOXN3 were replaced with the corresponding amino acids from FOXJ3, which is monospecific for FKH binding, and the complete sequence preference of the resulting chimeric domain was assayed *in vitro* in an unbiased manner. This analysis demonstrated that a portion of wing 2, which shows the largest difference in protein positioning between the FKH- and FHL-bound states (Figure 1C), is an important determinant of bispecificity. Newman *et al.* focused on the loop connecting helix 2 and helix 3 (hereafter referred to as the “2-3 loop”), demonstrating that several non-DNA-contacting positions within this loop vary systematically between the specificity classes of the FH domain (Figure 1D). They presented a crystal structure of the FHL-preferring FOXN1 protein bound to an FHL sequence and suggested that specific features of the 2-3 loop would constrain the ability of the recognition helix asparagine to adopt the FKH-selecting or FHL-selecting rotamers, but did not test this hypothesis by assaying the activity of chimeric proteins. Thus, the determinants of specificity within the FH family remain unresolved.

In addition to structural and biochemical studies focused on individual model proteins, such as described above for FOXN proteins ^24,25^, analysis of genetic variants can help to reveal amino acid residues important for protein function. We have previously shown that determining the effects of coding variation on the DNA-binding properties of human TF DBDs *in vitro* can provide insights into the amino acids that affect DNA sequence specificity, as well as clarify the mechanisms by which known disease-associated variants impact function and nominate additional variants present in the population as candidate functional alleles ^7,8^. We therefore performed a focused survey of FH domain missense variants and chimeric constructs to clarify several outstanding questions about the determinants of FH DNA binding bispecificity. Variants affecting FH family TF protein sequence have been identified in the human population; many of these have been associated with severe congenital diseases of the brain, lungs, lymphatic vessels, immune system, and other organ systems, from all germ layers ^26–31^. Others are Variants of Uncertain Significance (VUS) or entirely uncharacterized variants detected by whole-genome or whole-exome sequencing ^32^. Here, we tested 12 reference FH proteins and 61 FH missense variants identified in patients or in population surveys, in parallel with their reference protein alleles, by protein-binding microarrays (PBMs) to characterize the effects of each variant on its DNA binding activity. We also designed 5 additional single amino acid variants, and 17 more complex subdomain swap experiments guided by recently published structural insights, to further explore the determinants of binding specificity. We report, for the first time, a set of changes that can confer bispecificity on a formerly monospecific FH domain. Our results highlight the need to test variants at non-DNA-contacting amino acid residues for their potential effects on DNA binding activity.

## Results

### Prioritization of population variants for testing

We began with a list of 49 human FOX genes ^33^, and identified the position of the FH domain in the canonical isoform of each as defined by UniProt ^34^. We aligned all human FH domains and removed positions gapped in the majority of aligned domains to generate a common coordinate system for positions within the domain, which reproduced the positions in Pfam domain PF00250 ^35^. We filtered variants present in the ClinVar database of variants with interpretations of their clinical significance ^36^ and the Genome Aggregation Database (gnomAD) of human genetic variation from diverse populations ^32^ for single amino acid substitutions within the FH domain or within 15 positions upstream or downstream. We identified 2,402 such variants in gnomAD and 245 in ClinVar (Figure 2A,B, Table S1). Of the 245 ClinVar variants, 12 were classified as Benign or Likely Benign, 130 as Pathogenic, Likely Pathogenic, or Risk Factor, and 103 were of Uncertain significance or had Conflicting interpretations of pathogenicity (*i.e.*, VUS).

**Figure 2:**
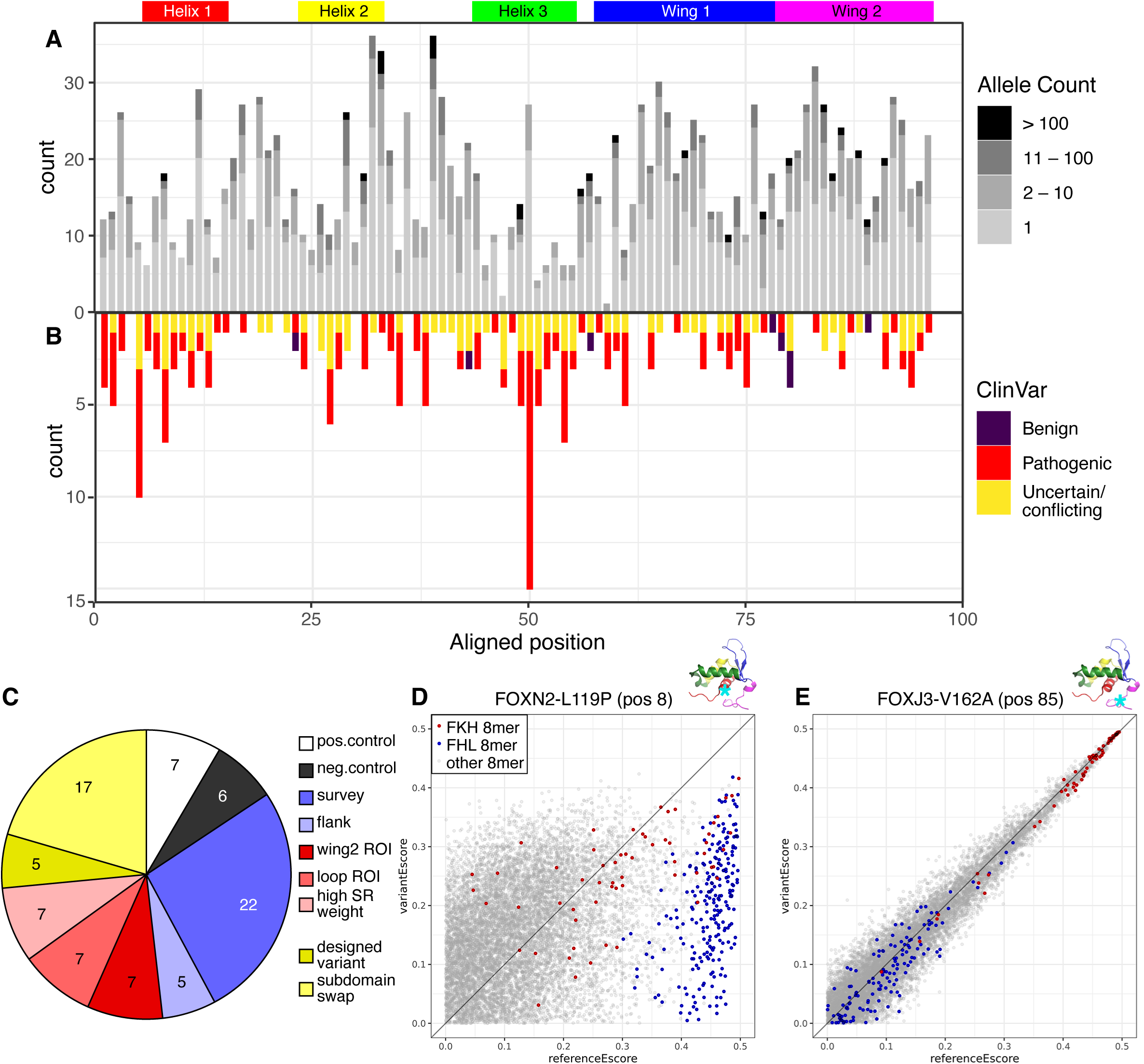
Identifying and testing FH domain variants present in the human population. **(A)** Missense variants are present in all regions of the FH domain in the gnomAD database (upper panel), usually at low frequency. **(B)** Missense variants in ClinVar are predominantly pathogenic and similarly distributed throughout the FH domain. **(C)** A total of 83 variant FH domains were tested in this study by PBMs to characterize their DNA binding preference, including 61 naturally occurring and 5 designed missense variants, and 17 more complex subdomain swaps. **(D)** The FOXN2 L119P variant was tested as a positive control for detection of disrupted binding. PBM enrichment (E) scores are shown for 8-mers of the TF reference allele on the x-axis (“referenceEscore”) and the variant on the y-axis (“variantEscore”). Points shown in red and blue represent 8-mers matching the FKH and FHL motifs, respectively. Points below the diagonal are 8-mers with reduced binding. In this and all subsequent figure panels, the position of the variant is shown on the FOXC2 structure for reference (cyan asterisk); note that the structure of wing 2 varies widely among FH subfamilies. **(E)** The FOXJ3 V162A variant, which is not predicted to affect binding, was tested as a negative control. E-scores are tightly clustered along the diagonal, indicating highly correlated binding by the variant and reference alleles. Scatterplots are representative of replicate PBM experiments.

We selected a range of variants to test for direct effects on DNA binding by performing protein binding microarray (PBM) experiments on both variant and reference alleles in parallel ^7,8^ (Figure 2C, Table S2, Figure S1). We selected several variants in the two regions previously highlighted as contributing to FH domain motif selectivity, the loop between helices 2 and 3, and wing 2. To narrow the large search space, we focused on variants predicted to be deleterious by all four of the popular variant interpretation tools SIFT ^37,38^, Mutation Assessor ^39^, PROVEAN ^40^ and PrimateAI ^41^. We tested several variants at positions with high Similarity Regression (SR) weight, a computational prediction of importance for sequence-specificity ^42^, again restricting to predicted deleterious variants. We also tested several predicted deleterious variants in positions flanking the rigorously defined FH domain, to assess the extent to which these protein regions must be taken into account when predicting effects on DNA binding. Finally, we tested a sample of variants at putative surface-exposed positions far from the DNA interface, surveying various regions of the FH domain. Non-DNA-contacting amino acid residues have been shown to influence the sequence specificity of several classes of TFs by various mechanisms ^8,9,43,44^, and specificity effects have proven difficult to predict *a priori* ^8^, motivating a survey approach.

As positive controls for our ability to detect effects on binding by this assay, we selected a number of variants which change an ultraconserved (across all FH subfamilies) DNA-contacting residue or insert a helix-breaking proline residue into one of the alpha helices that organize the domain. For example, FOXN2 L119P (position 8) inserts a proline into helix 1 and is predicted to be inconsistent with the correct folding of the domain. Scatterplots of the PBM enrichment (E) scores for all possible 8-mers depict the preference of a TF mutant allele (y-axis) versus the corresponding TF reference allele (x-axis) for binding 8-mers matching the FKH (red points) or FHL (blue points) motifs (Figure 2D,E); higher E-scores indicate 8-mers bound more preferentially. While FOXN2 can recognize both the FKH and FHL motifs, FOXN2 L119P causes an essentially complete loss of binding, with almost no 8-mers with E > 0.4 (representing strongly preferred binding), in 2 independent replicates (Figure 2D and Figure S1BD), and all 7 positive control variants tested similarly compromise DNA binding (Figure S1P,U,V,X,AP,BD,BY). As a negative control, we tested the FOXJ3 V162A (position 85) variant, which is present in the population at a high allele frequency (0.806). FOXJ3 reference and V162A variant E-scores were extremely well correlated in two replicate experiments (Figure 2E and Figure S1AW), consistent with no significant difference in their DNA binding activities. Other variants tested as negative controls were biochemically conservative changes at residues not predicted to contact DNA, and 0/6 negative control variants showed unambiguous effects on binding (Figure S1B,H,AJ,AN,AO,AW).

### Many gnomAD and ClinVar FH coding variants confer reduced DNA binding affinity

Several of the variants tested showed reduced binding activity (Table S3, Figure S1). For example, the likely pathogenic variant FOXF1 Y89C (position 42) has been associated with alveolar capillary dysplasia with pulmonary venous misalignment (ClinVar accession # SCV001427080) and showed severely reduced binding specificity for *k*-mers matching the FKH motif (Figure 3A). Some rare variants observed in the population but not associated with disease, *e.g.*, FOXB2 E52G (position 40), showed a similar decrease in binding (Figure 3B).

**Figure 3:**
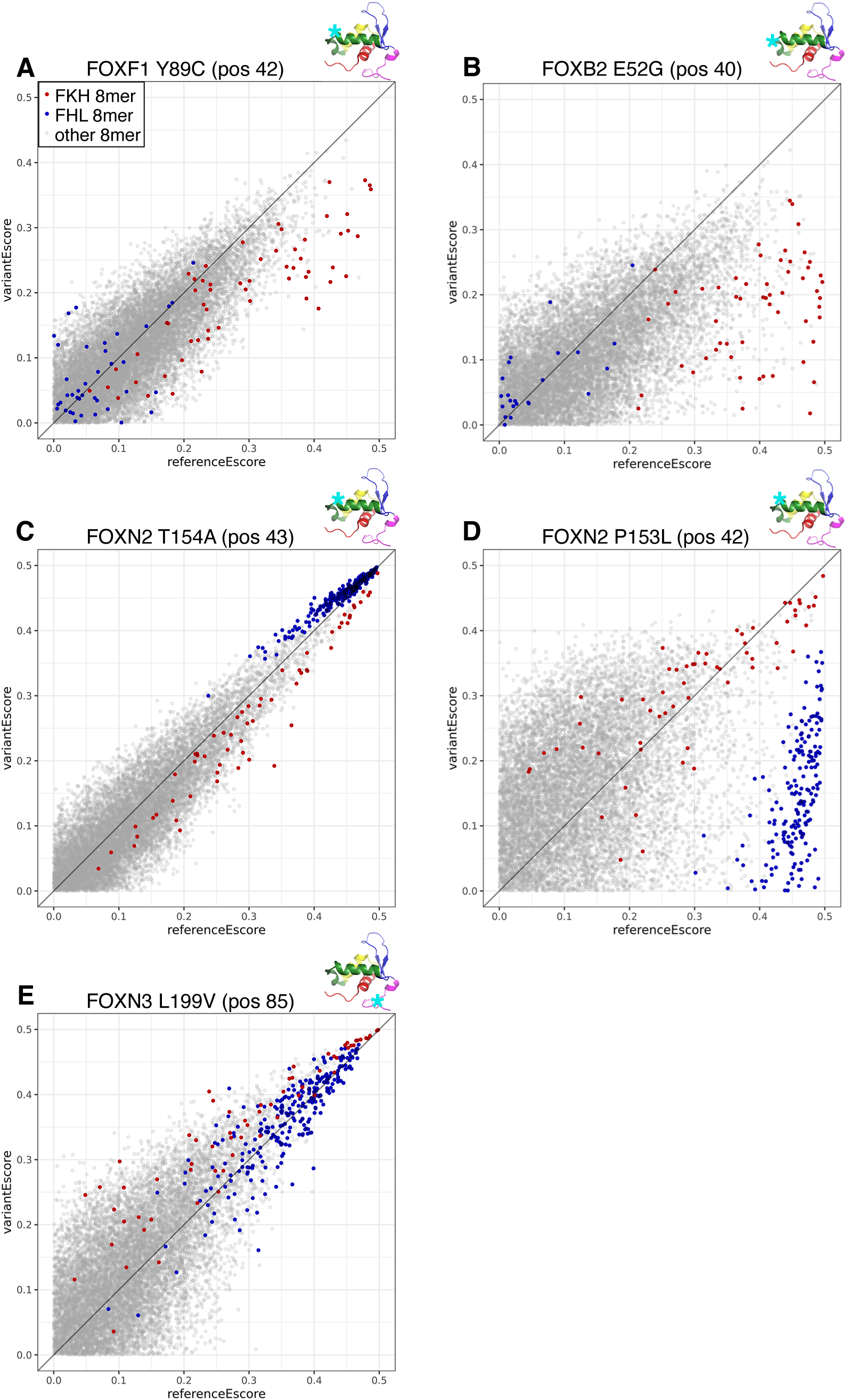
FH variants affecting DNA binding affinity or specificity. **(A)** FOXF1 Y89C is a likely pathogenic variant that causes a severe reduction in binding to FKH 8-mers (red points). **(B)** FOXB2 E52G is an uncharacterized rare variant that similarly reduces FKH motif binding. **(C)** FOXN2 T154A preserves binding to both FKH and FHL motifs, but alters specificity, with increased preference for FHL 8-mers (blue points) and decreased preference for FKH 8-mers (red points). **(D)** FOXN2 P153L severely impairs binding to FHL 8-mers while largely preserving FKH binding. **(E)** FOXN3 L199V preserves binding to both FKH and FHL motifs, but increases preference for FKH vs. FHL 8-mers. Scatterplots are formatted as in Figure 2 and are representative of replicate PBM experiments.

Variants that appear to impact binding were distributed throughout several locations distal to the DNA-binding surface, including the loops between the major helices and the wings (Figure S1I,O,AB,AM,CD). None of the flanking region variants we tested detectably impacted DNA binding (Figure S1F,N,S,AK,AR). A total of 19 naturally occurring variants in disease-associated genes showed alterations in binding; 16 of these variants had not been reported previously (Table 1).

**Table 1:**
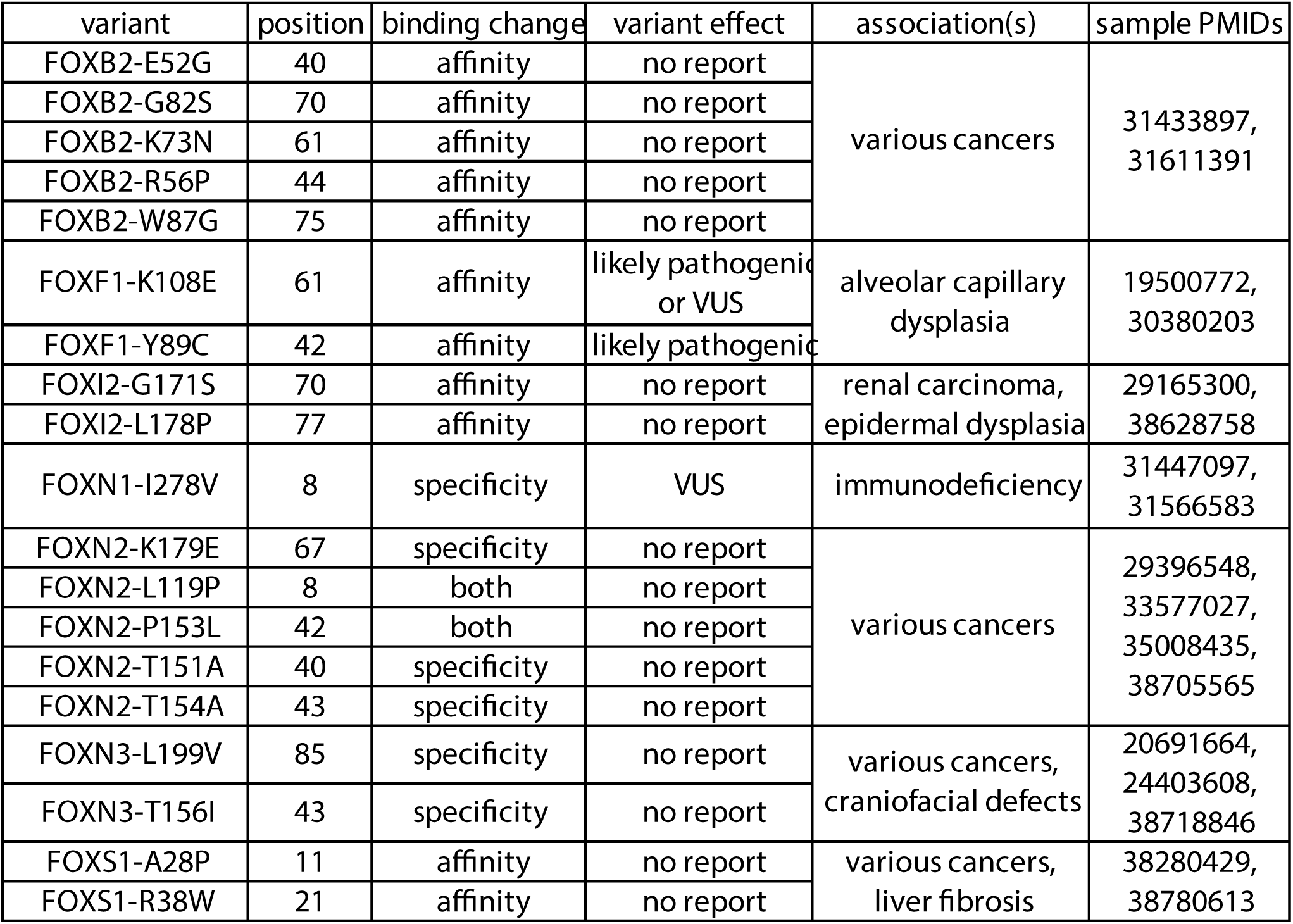
Altered DNA binding activities not previously identified for naturally occurring forkhead variants in disease-associated genes. . For each variant, we report its position within the domain, the observed DNA-binding change (affinity, specificity, or both), whether the variant has itself been reported to be pathogenic, the disease or diseases associated with the variant or gene, and selected references.

### gnomAD variants in FH wings and loop that support their roles in specificity

Several tested variants of bispecific forkheads present in the population affected the relative preference for FKH vs. FHL motifs. Two rare variants in the 2-3 loop of FOXN2 altered the preference in opposite directions. FOXN2 T154A (position 43) showed consistently increased preference for 8-mers matching the FHL motif, relative to 8-mers matching the FKH motif (Figure 3C). In contrast, FOXN2 P153L (position 42) largely eliminated preferential binding to FHL motifs, with selective sparing of FKH binding (Figure 3D). These findings are consistent with a prior prediction that the precise identities of amino acids at these positions are important for positioning the recognition helix asparagine residue that adopts consistently different rotamers between the FKH- and FHL-bound structures ^25^.

Newman *et al.* further speculated that the aspartic acid residue at position 43 in FOXN1 may contribute to the strong FHL preference that we previously characterized as monospecificity of that factor ^25^, by shifting the 2-3 loop to avoid an unfavorable charge interaction with the DNA backbone. We therefore tested a designed mutation at that position in a bispecific forkhead, FOXN3 T156D (position 43). We observed a subtle shift toward FHL binding by this mutant, but FKH binding was strongly maintained (Figure S1BQ). Similarly, we tested the gnomAD variant FOXN1-D313N at this position and observed no consistent effect on binding or specificity (Figure S1AX). In particular, we detected no increased binding to FKH 8mers. An additional rare variant present in the population, FOXN3 L199V (position 85), maps to the region of wing 2 previously implicated in FKH vs. FHL specificity ^24^. This very conservative amino acid change caused a subtle but highly consistent shift toward FKH vs. FHL preference (Figure 3E and Figure S1BL), consistent with a role for wing 2 in determining specificity.

### FKH vs. FHL preference of bispecific factors is likely relevant *in vivo*

To address the question of whether these population variants were likely to have phenotypic effects, we analyzed published ChIP-seq data for bispecific FOXN3 from HepG2 and MCF-7 human cell lines ^24,45^. FOXN3 is a transcriptional repressor [16102918] that has been implicated in human craniofacial defects ^46^ and various cancers ^47,48^. Both the FKH and FHL motifs are enriched among ChIP-Seq peaks in at least one of these two datasets, as compared to a matched genomic background (Figure 4A,B; see Methods). Using the GREAT tool ^49^ to associate ChIP peaks with genes and enriched gene sets, the subset of peaks containing matches to the FKH motif had no significantly enriched Gene Ontology (GO) or phenotype associations in common between the two datasets. Therefore, we focused subsequent analyses on each cell line individually.

**Figure 4:**
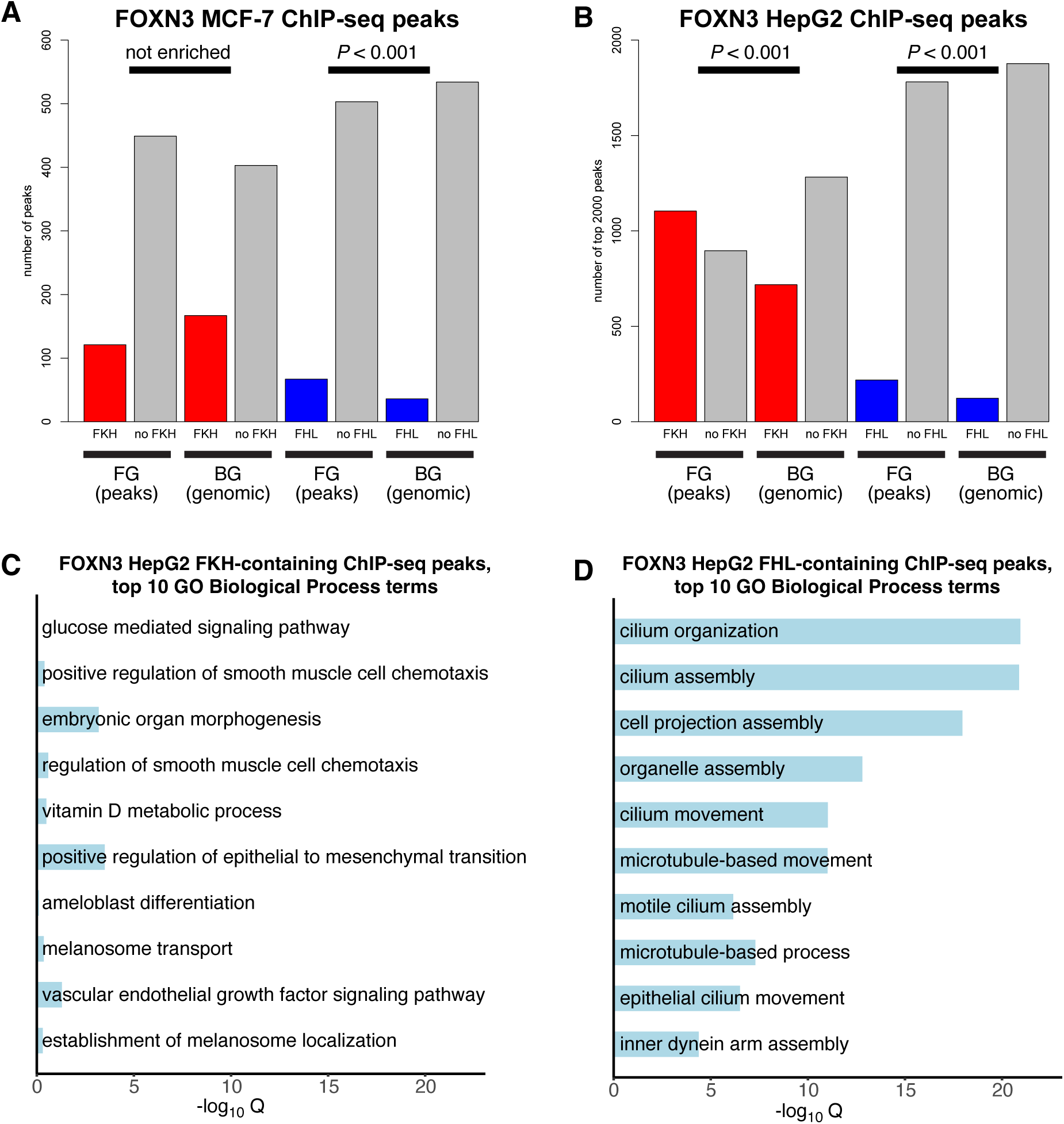
FHL binding mode of FOXN3 is associated with regulation of cilia-related genes. **(A)** FOXN3 ChIP-Seq peaks in published data from human MCF-7 cells show enrichment of FHL 8-mers (blue bars) vs. matched genomic background, but not of FKH 8-mers (red bars). *P*-values from Fisher’s Exact Test. **(B)** FOXN3 ChIP-Seq peaks in published data from human HepG2 cells show enrichment for both FKH and FHL 8-mers vs. matched genomic background. *P*-values from Fisher’s Exact Test. **(C)** Top 10 GO Biological Process terms enriched in targets of FKH-containing FOXN3 peaks in HepG2 cells. GREAT calculates both enrichment of regulatory regions (with significance calculated by binomial test) and of putative target genes (with significance calculated by hypergeometric test); the categories displayed here and panel (D) are ranked by binomial enrichment in regulatory regions and significance shown is for target gene enrichment. Enriched terms show little significant target gene enrichment and no common theme. **(D)** Top 10 GO Biological Process terms enriched in targets of FHL-containing FOXN3 peaks in HepG2 cells; genes involved in cilia-related processes are much more enriched among these target gene sets. Category rankings and significance shown are as in panel (D).

Focusing on the FOXN3 ChIP-Seq peaks in HepG2 cells, the top 10 Biological Process terms in peaks containing matches to the FKH motif similarly showed no common theme and very little statistical significance (Figure 4C). The peaks containing matches to the FHL motif, in contrast, had 31 GO terms significantly enriched in both datasets. These enriched GO terms all concerned the cilium and its assembly and organization. 8 significantly enriched human phenotype associations in common were all explainable by ciliary dysfunction, as illustrated by the top 10 terms enriched in HepG2 FHL ChIP-Seq peaks (Figure 4D). Results for FOXN3 ChIP-Seq peaks in MCF-7 cells were similar: 7 of the top 10 enriched GO terms among FHL peaks were also among the top 10 terms in HepG2 cells, while no terms were significantly enriched among FKH peaks and none of the top 10 terms enriched among FKH peaks were common between the two cell types. This suggests that targeting of the cilium assembly pathway by FOXN3 could be mediated preferentially by recognition of the FHL motif, and therefore that variants changing the relative preference of FOXN3 for FKH vs. FHL targets could alter downstream regulatory effects. Moreover, targeted disruption of the *Foxn3* gene caused dysregulation of cilia-associated genes in the embryonic mouse retina, along with disorganized and abnormal cilia ^50^. Altogether, these results suggest that the FHL binding mode is used by FOXN3 to regulate cilia-associated genes.

### Subdomain swaps support a distributed model of FH specificity

The observations that no tested missense variant switched the specificity of a monospecific forkhead from FKH to FHL or vice versa or to bispecificity, and that variants affecting the relative preference of bispecific forkheads for FKH vs. FHL motifs cluster in two parts of the domain, suggest a model in which both subdomains must be permissive for binding a given motif. As in previous experiments ^24^, we tested this idea by engineering “subdomain swaps,” in which larger portions of one FH domain are replaced with the aligned portion of another. Consistent with this model, replacing the 2-3 loop of bispecific FOXN3 with that from FKH-monospecific FOXA1 abrogated FHL recognition without compromising FKH binding (Figure 5A), while replacing the wings of FOXA1 with those of FOXN3 spares FKH binding but is unable to confer FHL specificity (Figure 5B). Similarly, replacing the 2-3 loop of FOXN3 with that from FHL-preferring FOXN1 causes a slight but consistent increase in specificity for the FHL vs. FKH motif (Figure 5C), while the reciprocal replacement does not render FOXN1 bispecific (Figure 5D), presumably due to a requirement for an FKH-binding wing configuration.

**Figure 5:**
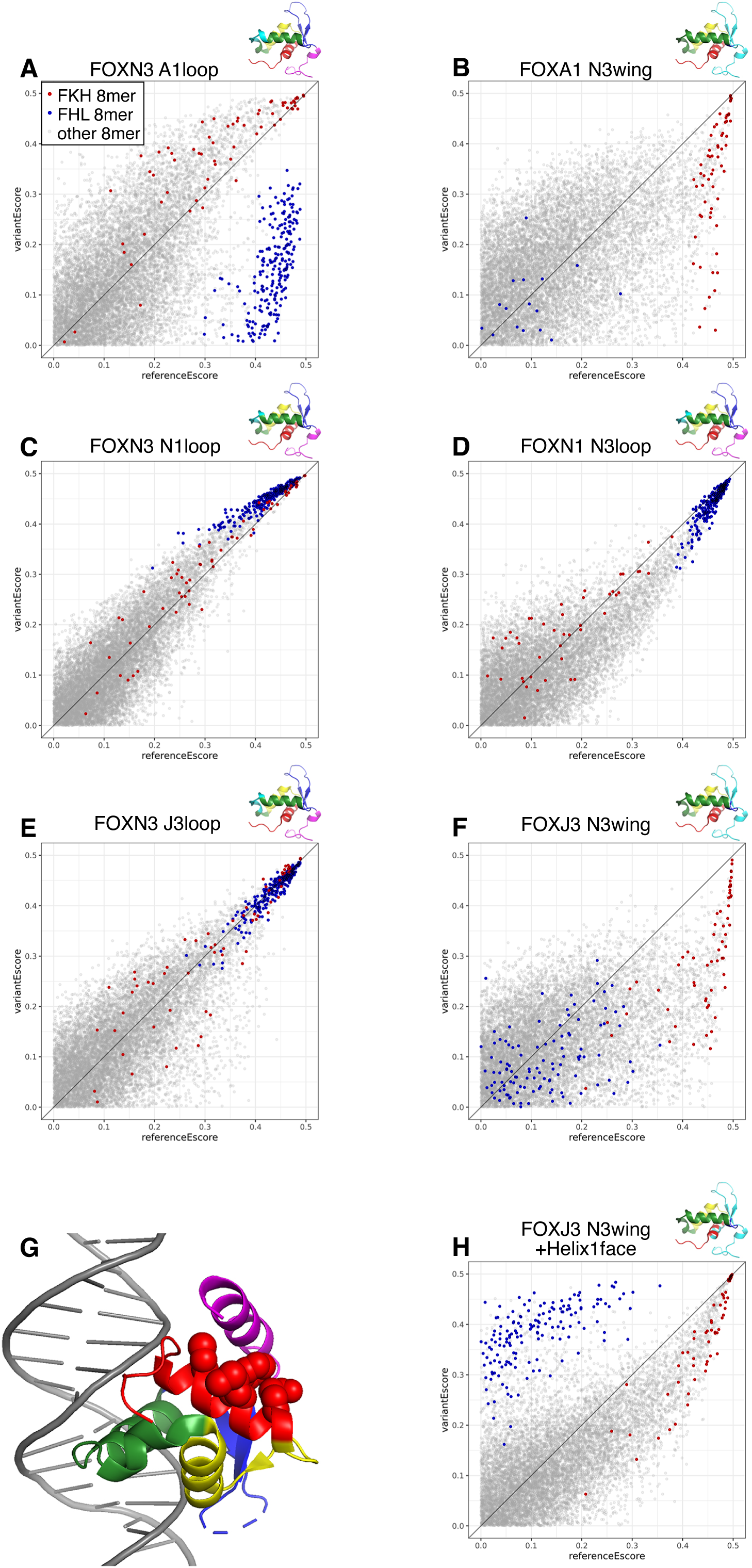
Determinants of FH bispecificity. **(A)** Replacing the loop between helix 2 and helix 3 (2-3 loop) in FOXN3 with the corresponding amino acids from FOXA1 abrogates FHL binding while preserving binding to FKH-containing 8-mers. In this and all subsequent panels, the region shown in cyan in the small schematic is the portion of the domain that differs from the reference allele, shown on the FOXC2 structure for reference; note that the structure of wing 2 varies widely among FH subfamilies. **(B)** Replacing the wings of FOXA1 with those of FOXN3 reduces FKH binding without conferring specificity for the FHL motif. **(C)** Replacing the 2-3 loop of FOXN3 with that of FOXN1 shifts binding specificity toward FHL 8-mers. **(D)** The 2-3 loop of FOXN3 does not confer FKH binding capacity on FOXN1. **(E)** The 2-3 loop from FOXJ3 (FKH monospecific) does not compromise FHL binding in FOXN3 (data from ^24^). **(F)** The wings of FOXN3 are not sufficient to confer FHL binding on FOXJ3. **(G)** Wing 2 of FOXN3 packs against a contact surface formed by a face of helix 1 (side chains of these helix 1 residues are shown as red spheres). **(H)** Mutation of these contact residues to those present in FOXN3 permits the FOXN3 wings to confer FHL binding on FOXJ3. Scatterplots are formatted as in Figure 2 and are representative of replicate PBM experiments.

We have previously shown, however, that replacing the 2-3 loop of FOXN3 with that of FKH-monospecific FOXJ3 does not abrogate FHL binding (Figure 5E), implying that if our model is correct, then FOXJ3 has a 2-3 loop consistent with bispecificity. Nevertheless, when the wings of FOXJ3 were replaced with those from FOXN3, FHL binding was not conferred (Figure 5F). Subsequently, a study of disease mutations in FOXG1 noted that Pfam position 95 within wing 2 appears to destabilize the domain when mutated, reducing its binding to a high affinity FKH site, that several wing 2 residues (primarily positions 81 and 84) make intramolecular contacts with helix 1, and that these contacts may be conserved in other FH domains ^23^. We therefore inspected the intradomain contacts made by wing 2 in different published FH structures ^21,22,24,25,51^ and found that, while the identities of the amino acids at the involved positions vary between family members, there is a conserved contact surface between wing 2 and a face of helix 1 (Fig. 5G). Moreover, wing 2 of FOXN3 packs against helix 1 in this way (Figure 5G), and there are several positions (Pfam positions 8, 11, 12, and 15) within this interface surface of helix 1 that are not conserved in FOXJ3, implying that the FOXN3 wings would not properly pack against the FOXJ3 helical bundle. We thus replaced an additional 4 amino acids in helix 1 of the FOXJ3-N3wing construct with the corresponding residues from FOXN3. This reproducibly conferred bispecificity on the previously FKH-monospecific FOXJ3- N3wing construct (Figure 5H), demonstrating that in the presence of a permissive 2-3 loop, a properly folded wing 2 can support binding to both the FKH and FHL motifs.

## Discussion

Nonsynonymous variants in TF DBDs are prevalent in the human population ^7^, and many of these variants are associated with Mendelian disease or with heightened disease risk ^36,52^. In this present study, we have identified thousands of such variants in the Forkhead family of TFs. However, not all variants will affect the DNA-binding activities of these proteins, and testing of individual variants is labor-intensive, necessitating a strategy for prioritizing those variants likely to be functionally relevant. Variants which affect the affinity of DNA binding can preferentially alter binding to lower affinity binding sites ^8^, and these moderate affinity motif instances have been shown to be crucial for the precise and flexible regulation of gene expression *in vivo* both in human disease and in developing model organisms ^53–57^. Reduced binding affinity, by reducing total TF occupancy, is analogous to a reduction of TF expression level; a subset of *in vivo* targets have been shown to be preferentially sensitive to the level of SOX9 expression, explaining features of the syndrome observed in haploinsufficiency ^58^.

Alteration of TF DNA-binding specificity can also cause human disease ^7,59,60^. Here, we present evidence that the bispecific FOXN3 TF may regulate the transcriptional program for cilia formation preferentially through the recognition of FHL motifs in the genome, and *in vivo* targets with matches to the FHL motif are enriched for genes associated with ciliopathies, suggesting that variants which alter this specificity could be significant drivers of disease in ciliated cell types. Existing computational tools for variant effect prediction struggle to capture the likelihood that a variant will affect the binding activity and, particularly, the sequence specificity of a TF ^8^. Understanding the determinants of binding affinity and specificity within each TF class will therefore be necessary to interpret the increasing amount of genomic sequence data now becoming available.

Here, we have shown that the wings and the 2-3 loop of a FH domain are together sufficient to determine preferential binding to the FKH or FHL motif, or bispecificity. This restricts the potential set of variants which are candidates for specificity altering. However, it remains unknown exactly what constitutes a permissive wing or loop for each motif. As discussed above, we observed no effects consistent with an aspartate residue at position 43 selecting FHL specificity. Similarly, we tested the ability of the FOXO3 loop to support bispecificity when provided with the wings of bispecific FOXN3, because the FOXO3 loop is rich in small, flexible amino acids, which have been hypothesized to support FHL binding ^25^, but found no effect on specificity (Figure S1BU-BW). The threonine residue at position 43 in bispecific forkheads that does affect binding when mutated does not contact DNA in published crystal structures bound to either motif ^24^. One possibility is that these residues near—but not touching—the protein-DNA contact surface may affect the network of ordered water molecules present in and around the interface. FH domain DNA recognition has been reported to rely on a unique network of water interactions ^15,61^, and water-mediated contacts have been implicated in position-interdependent recognition of nucleotide combinations by other classes of TF ^62^. Higher-resolution structural studies of reference and variant alleles bound to DNA will be necessary to fully understand the features that determine specificity. This is especially relevant for the FH domains because there have been reports of surprising additional specificities in this family, including variations on the FHL motif in FOXM1 ^20^, FHL recognition as a half-site in the context of dimer binding to an FKH site by FOXP3 ^63^ and a novel AATCCACA motif recognized by FOXH1 ^64^.

Studies of TF-DNA binding specificity typically have focused on DNA-contacting residues. The results presented here reveal amino acid residues, located in regions of FH domains that do not contact DNA that allosterically and synergistically control whether a FH TF will be monospecific for either the FKH or FHL motif, or bispecific for both motifs. Recognition of multiple DNA sequence motifs has been reported for TFs from other structural classes ^65–69^.

Non-DNA-contacting amino acid residues of a broader set of TFs may contribute to the relative preferences among their recognition motifs and may play a role in the evolution of TF DNA binding specificities and gene regulatory networks.

## METHODS

### Forkhead amino acid sequence alignment

We downloaded the hidden Markov model (HMM) representation of the FH domain (PF00250) from the Pfam database ^35^. We then searched the canonical amino acid sequences from UniProt ^34^ corresponding to the set of all human TFs, as defined by Lambert *et al.* ^33^, with the HMMER software suite ^70^ for matches to this pattern, and compiled a list of coordinates for the FH domain within the canonical protein product of each human gene. These domains were aligned with MAFFT ^71^ using the L-INS-i parameter set. The resulting alignment was consistent with the position numbering of the Pfam domain model after removal of highly gapped, subfamily-specific positions; this trimmed alignment was used to assign variants in human proteins to positions within the domain.

### Cloning of reference and variant forkheads

Forkhead reference and domain swap DBD regions, flanked by 15 additional amino acids N- and C- terminal of DBD and by Gateway attB recombination sites, were manufactured through gene synthesis (IDT gBlocks). Synthesized fragments were cloned into the Gateway compatible pDONR221 Entry clone and then transferred into pDEST15, an N-terminal Glutathione S-transferase (GST) protein fusion Destination vector. All cloned sequences were sequence confirmed by Sanger sequencing (Table S3).

Variant DBD clones were generated using site-directed mutagenesis. Site-directed PCR was performed using variant-specific, complementary forward and reverse primers (Table S3) on the DBD cloned into the pDONR221 Entry clone. Nine variants (FOXA1-4aa, FOXA1- N3wing-4aa, FOXJ3-4aa, FOXJ3-N3wing-4aa, FOXN3-J3wing-4aa, FOXO3-4aa, FOXO3-N3wing-4aa, FOXS1-4aa, and FOXS1-N3wing-4aa) required multiple rounds of site-directed PCR to generate all desired mutations. Mutagenesis clones were sequence-verified by Sanger sequencing before the variant DBD was then transferred into pDEST15.

### Protein expression

Reference and variant N-terminal GST-fusion proteins were expressed using the bacterial-based PURExpress *in vitro* transcription and translation (IVT) kit (NEB) according to the manufacturer’s protocol. Protein expression was confirmed, and molar concentrations were quantified by Western blot using a serial dilution of recombinant GST protein (Sigma) as standards. Equal volumes of expressed protein and GST of known molar concentrations were loaded and run on a NuPAGE 4-12% Bis-Tris gel (Invitrogen) and then transferred to a nitrocellulose membrane (Invitrogen). Membranes were blotted with a 1:200,000 dilution of rabbit anti-GST primary antibody (Sigma) followed by incubation with a 1:160,000 dilution of goat anti-rabbit secondary antibody (Invitrogen). Reference and variant alleles within an allelic series were expressed in the same IVT batch and quantified by Western blotting together.

### Protein binding microarray experiments

PBM experiments were performed using an “all 10-mer” universal array in 8 x 60K, GSE format (Agilent Technologies; AMADID #030236). Arrays were double-stranded and PBM experiments were completed following previously described protocols ^72,73^. All Forkhead TFs were assayed at a final protein concentration of 750 nM. All experimental comparisons between reference and variants were performed in separate chambers on the same slide. All arrays were scanned using a GenePix 4400A microarray scanner. For all PBM experiments, the array was scanned 3 times at different photomultiplier tube (PMT) gain settings. Protein concentration, buffer, and array ID of each PBM experiment are provided in Table S2.

### Statistical analysis of PBM data

PBM scans were analyzed using GenePix software and the Universal PBM Analysis Suite, as described previously ^73^. This yields an enrichment score (E-score) for every possible 8-mer in every replicate of every TF variant or its matched reference experiment performed in parallel (Table S4). To call changes in specificity, variant E-score minus reference E-score was calculated for all 8-mers matching either the FKH consensus (RYAAAYA) or the FHL motif (GACGC) and with an E-score ≥ 0.4 in either condition. The distribution of these difference scores was then compared for FKH-containing vs. FHL-containing 8-mers using the wilcox.test function in R. A specificity difference was called if these distributions differed in the same direction with an adjusted p-value (False Discovery Rate (FDR)) ≤ 0.1 for all replicates of a given variant. To call changes in affinity, we similarly applied the paired Wilcoxon test to variant vs. reference E-scores for the same subsets of 8-mers, but this analysis proved to be poorly calibrated when tested on reference-vs-reference control comparisons. We therefore performed this test on a large panel of 320 reference-vs-reference comparisons and selected an effect size cutoff representing a 10% empirical FDR. A variant was considered to affect binding affinity if the absolute value of its effect size exceeded this cutoff consistently in all replicates. In cases where replicates disagreed, we performed at least 1 additional replicate. We reasoned that a microarray experiment may fail to show specific binding for technical reasons, whereas it’s highly unlikely that a failed experiment would produce E-scores well correlated with a successful reference experiment performed in parallel; therefore, when 1 out of 3 replicates showed loss of specific binding and the remaining 2 did not, we considered this variant to not affect affinity. The resulting affinity and specificity calls are reported in Table S3.

### Statistical analysis of published data

For FOXN3 ChIP-Seq data from HepG2 cells, the top 2000 peaks were downloaded from Rogers et al. Table S1 ^24^. For MCF-7 cells, the raw reads from FOXN3 ChIP-Seq and Input samples were downloaded from GEO Series GSE93780 ^45^ and mapped to GRCh38 using Bowtie 2, and peaks were called using MACS2 with Input as background and default parameters. Each peak set was trimmed to 150 bp segments centered on peak summit, and random background sets matched for G/C content, CpG content, promoter overlap, and repeat content were generated using GENRE ^74^. Sequences containing matches to the FKH (RYAAAYA) or FHL (GACGC) motif were counted in each peak set and its respective background set, and statistical significance of enrichment was assessed with Fisher’s Exact Test in R. GO terms and phenotype associations enriched in sets of peak sequences containing a given motif match were assessed using GREAT version 4 ^49,75^.

## Data Availability

PBM data have been deposited in the GEO database under accession number GSE297784.

## Supplemental Information

Supplemental Information includes 1 figure and 4 tables.

## Acknowledgements

We thank members of the Bulyk lab, in particular Kaia Mattioli and Luca Mariani for helpful discussions and Julia Rogers for critical reading of the manuscript. This work was supported by the National Institutes of Health (grant R01 HG010501 to M.L.B.).

## Author Contributions

M.L.B. conceived the research project. M.L.B. and S.S.G. designed the research project. J.K. and E.R. performed experiments. J.K., S.S.G., J.-A.D., and R.J. performed analyses. S.S.G. prepared the figures. M.L.B. supervised the research. S.S.G. and M.L.B. wrote the manuscript. All authors approved the final version of the manuscript.

## Supplemental information

Figure S1 Tables S1-S4

## Supplementary Tables

**Table S1:** Single amino acid variants affecting the FH domain or flanks in gnomAD and ClinVar.

**Table S2:** Forkhead reference, variant or designed mutant proteins analyzed in this study, including clone sequences (DNA and a.a. sequence) and sequences of primers used in site-directed mutagenesis.

**Table S3:** PBM experiment conditions and result “calls” of altered DNA binding activity.

**Table S4:** All PBM E-scores for all 8-mers for all proteins analyzed in this study.

