## Supplementary material for "Missense variants in human forkhead transcription factors reveal determinants of forkhead DNA bispecificity": Figure S1

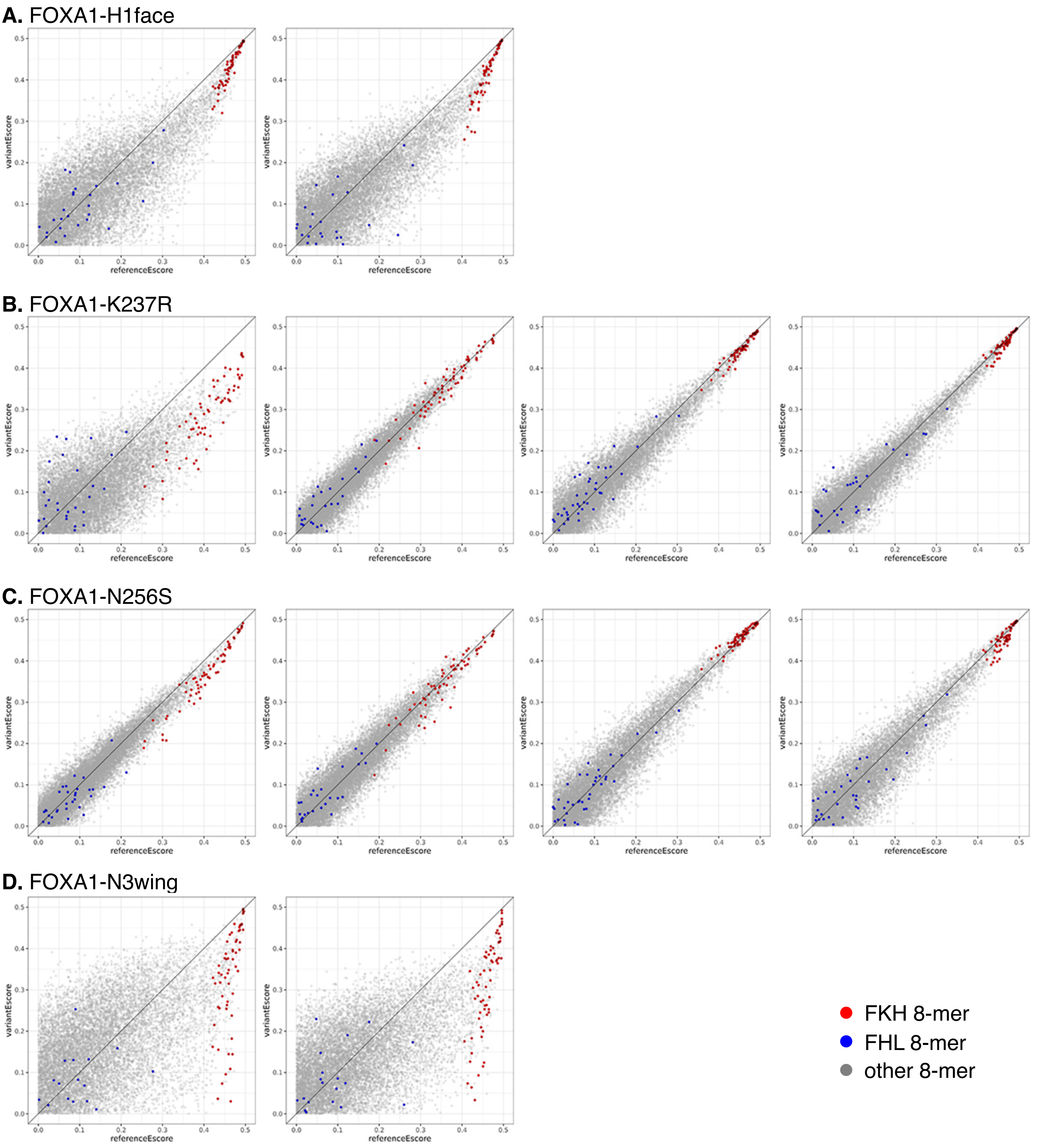


### Supplementary Figure 1

PBM enrichment (E) scores are shown for 8-mers of the reference allele on the x-axis and the variant on the y-axis, for all replicates performed. Points shown in red and blue represent 8-mers matching the FKH and FHL motifs, respectively.

**E.** FOXA1-N3wing+H1face

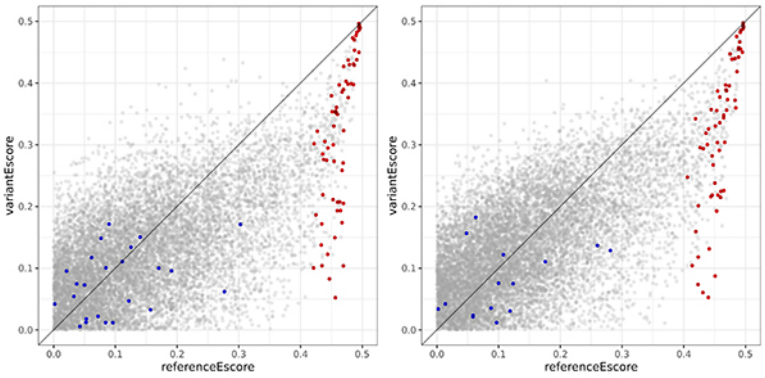


**F.** FOXA1-S165R

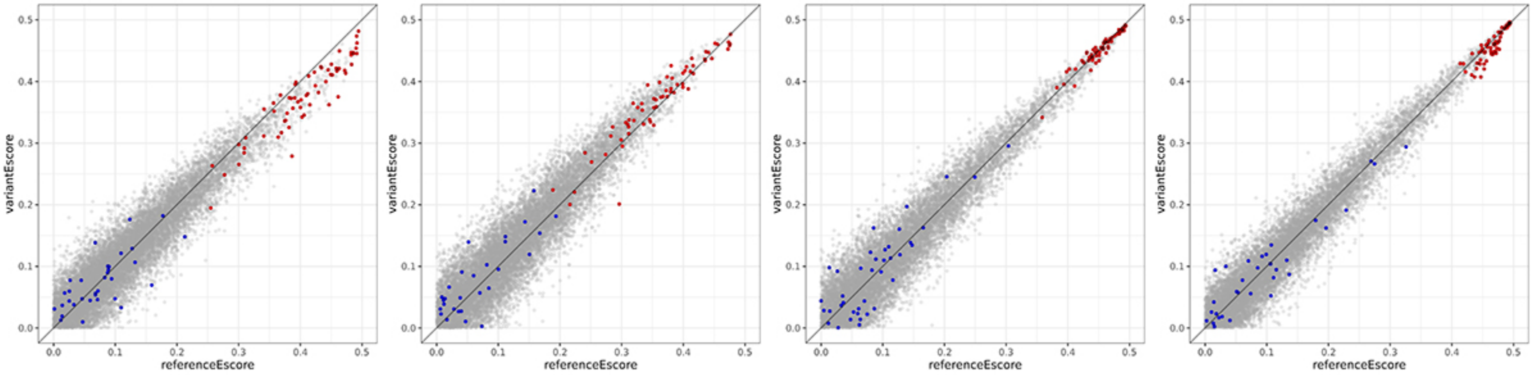


**G.** FOXA1-S188G

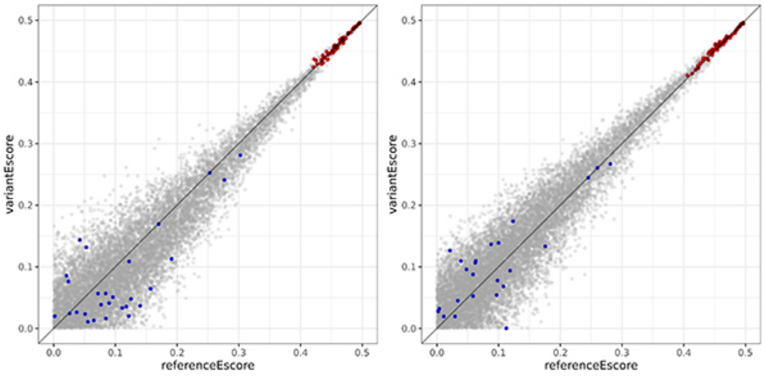


**H.** FOXB2-D95E

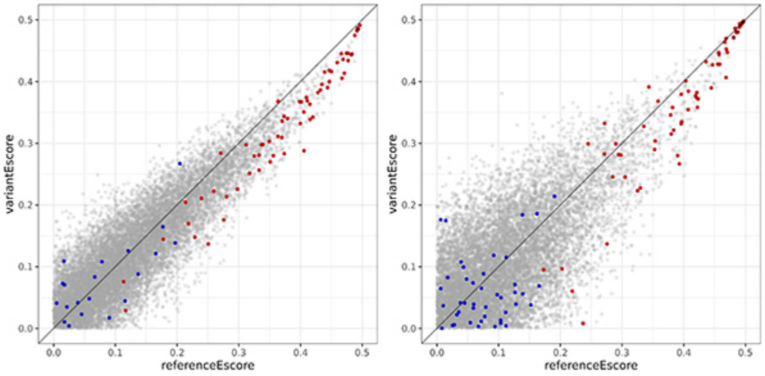


FKH 8-mer
FHL 8-mer
other 8-mer

**Supplementary Figure 1 (cont.)**

**I.** FOXB2-E52G

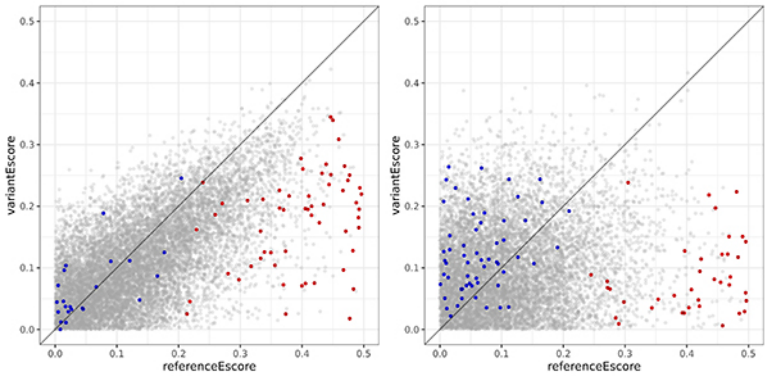


**J.** FOXB2-G100C

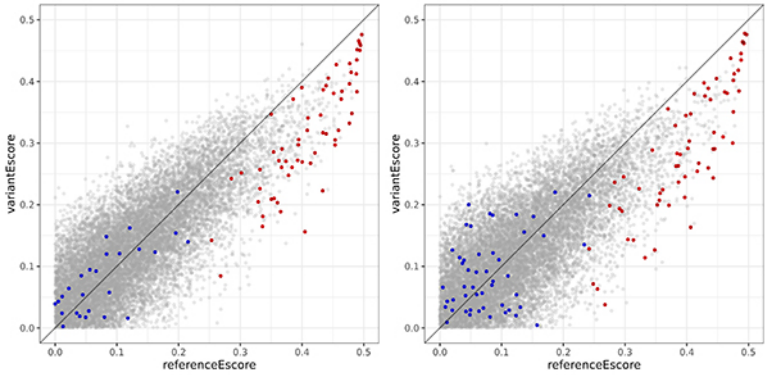


**K.** FOXB2-G100D

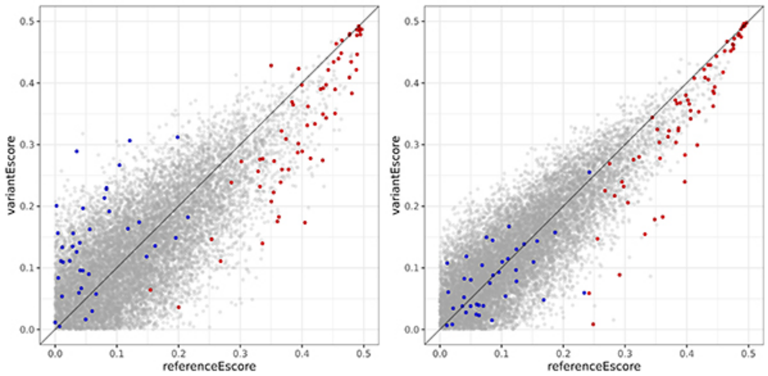


**L.** FOXB2-G100R

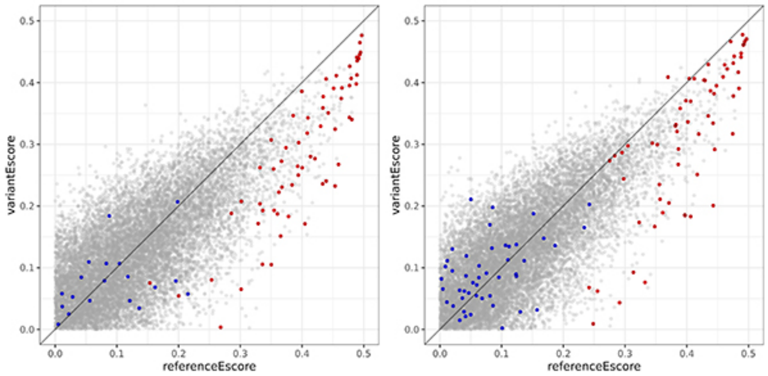


FKH 8-mer
FHL 8-mer
other 8-mer

**Supplementary Figure 1 (cont.)**

**M.** FOXB2-G100S

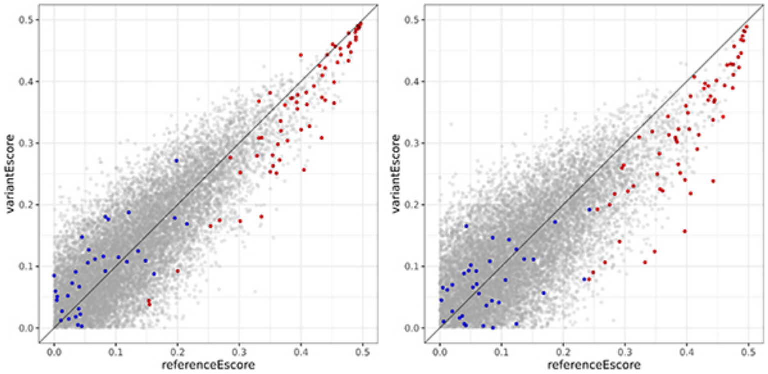


**N.** FOXB2-G5R

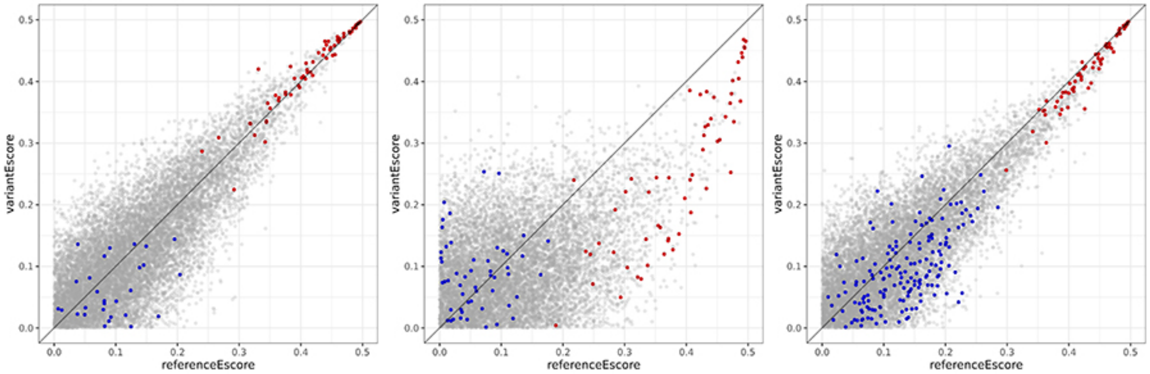


**O.** FOXB2-G82S

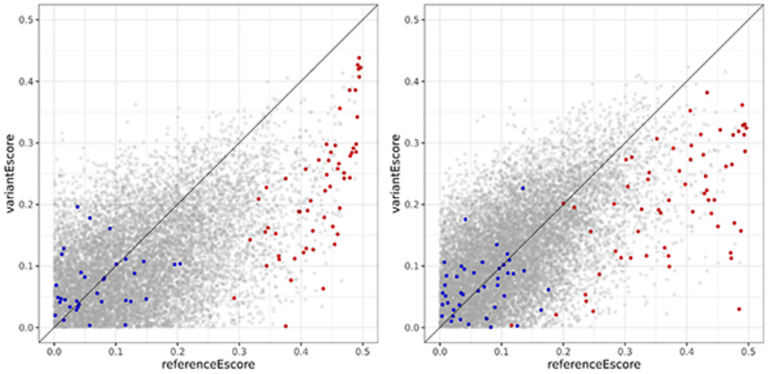


**P.** FOXB2-K73N

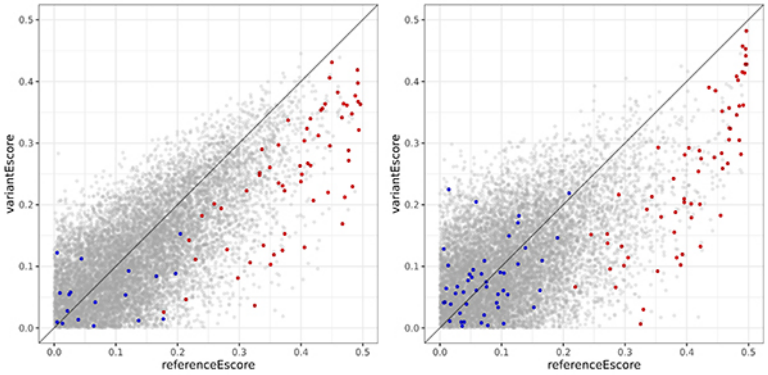


FKH 8-mer
FHL 8-mer
other 8-mer

**Supplementary Figure 1 (cont.)**

**Q.** FOXB2-P35L

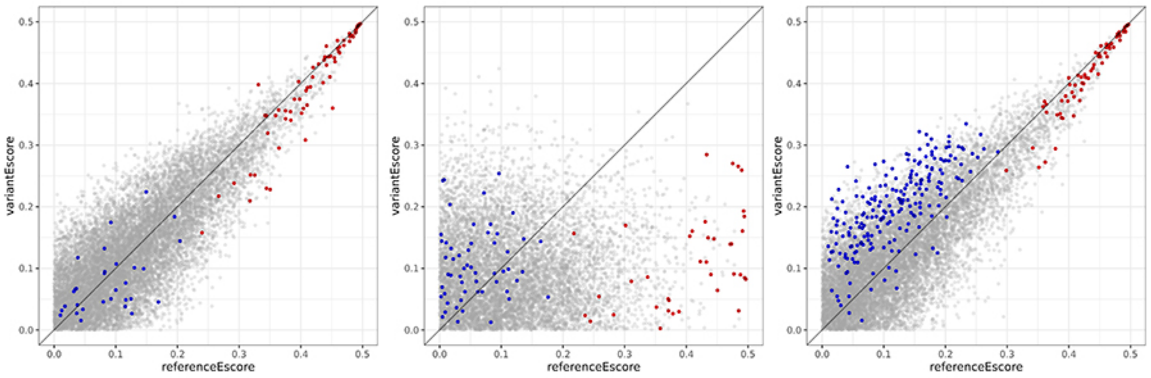


**R.** FOXB2-P35Q

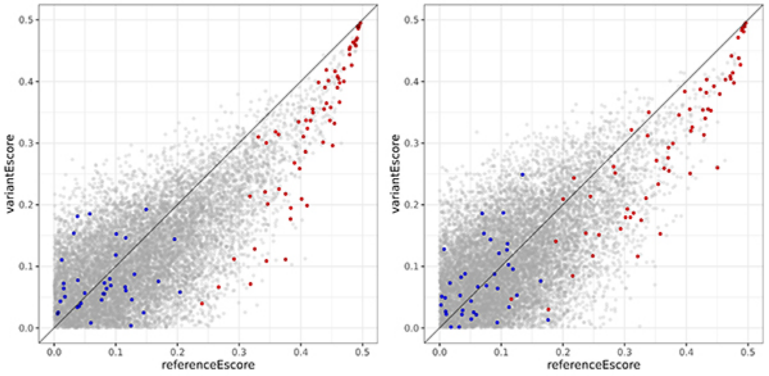


**S.** FOXB2-R3Q

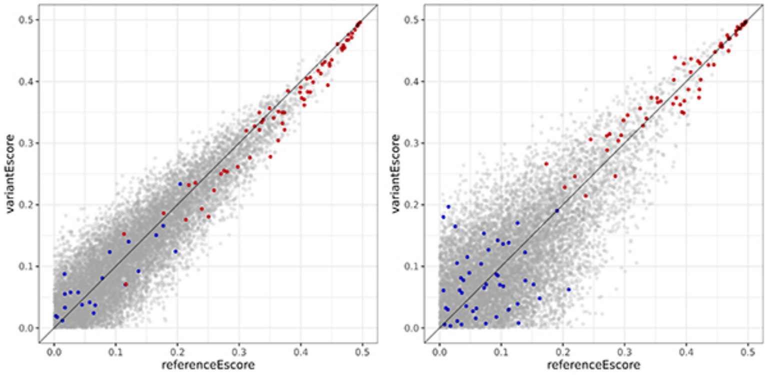


**T.** FOXB2-R46C

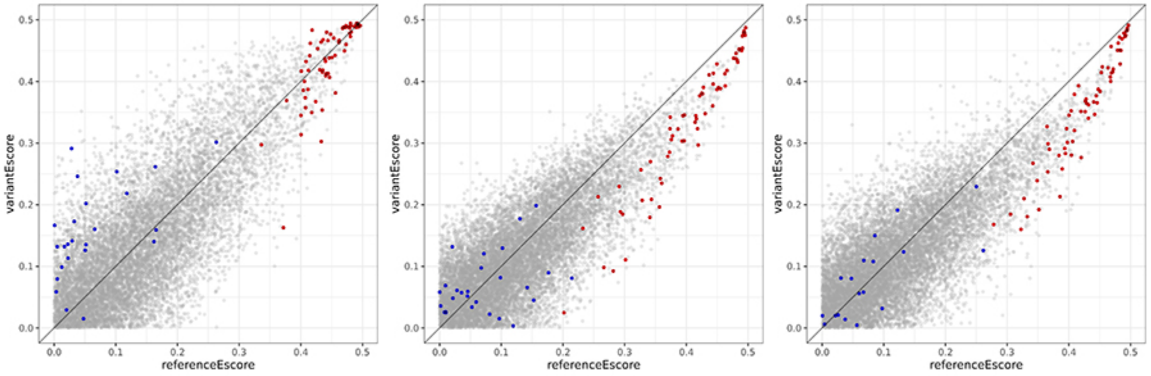


● FKH 8-mer
● FHL 8-mer
● other 8-mer

**Supplementary Figure 1 (cont.)**

**U.** FOXB2-R56P

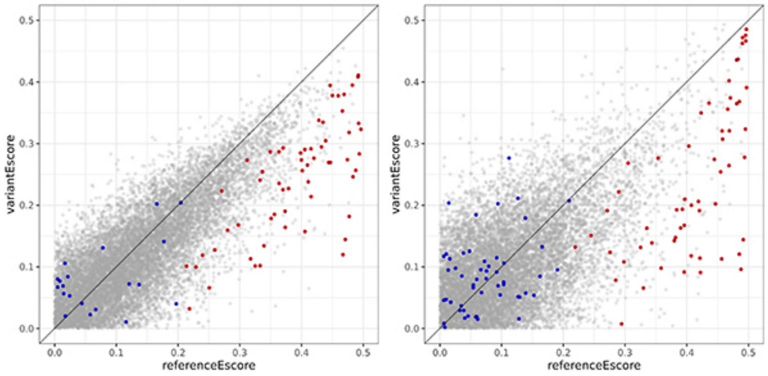


**V.** FOXB2-W87G

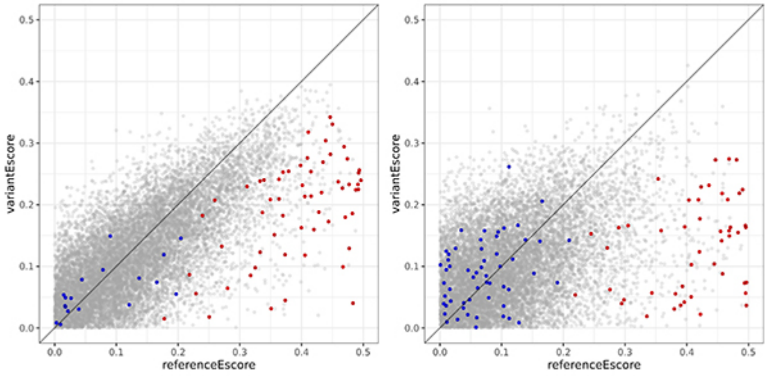


**W.** FOXF1-G112S

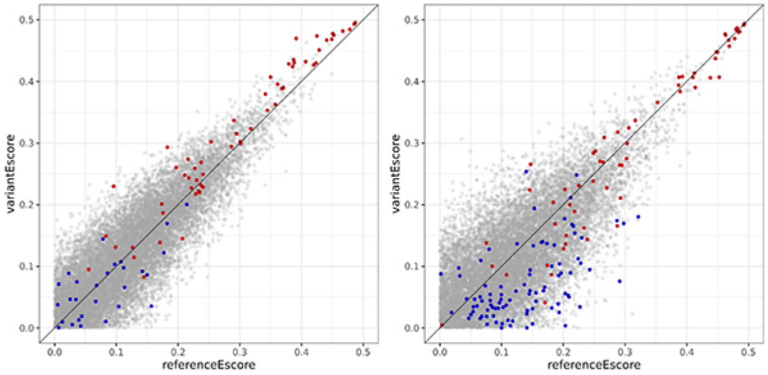


**X.** FOXF1-K108E

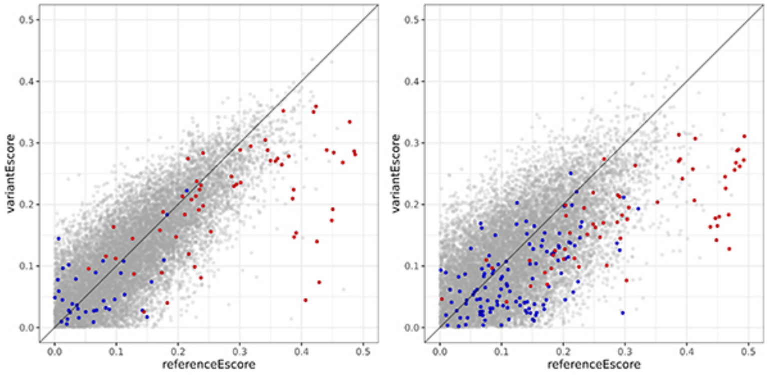


● FKH 8-mer
● FHL 8-mer
● other 8-mer

**Supplementary Figure 1 (cont.)**

**Y.** FOXF1-P83A

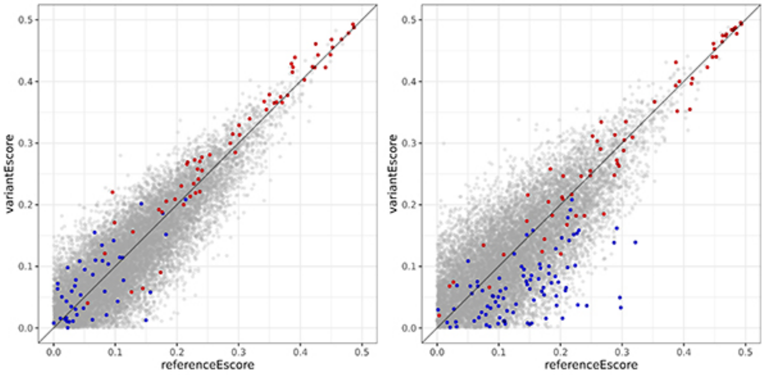


**Z.** FOXF1-P83S

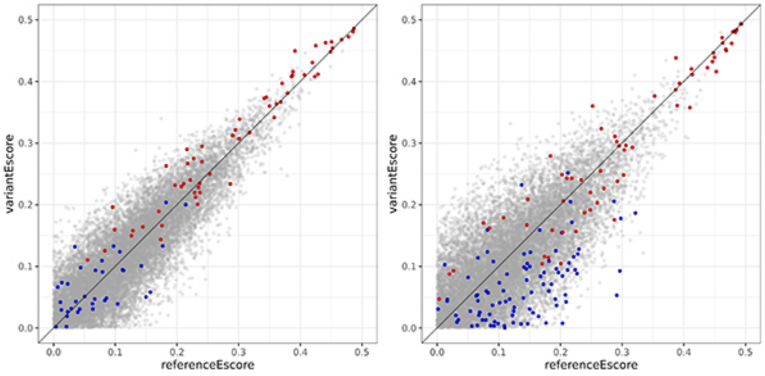


**AA.** FOXF1-R139Q

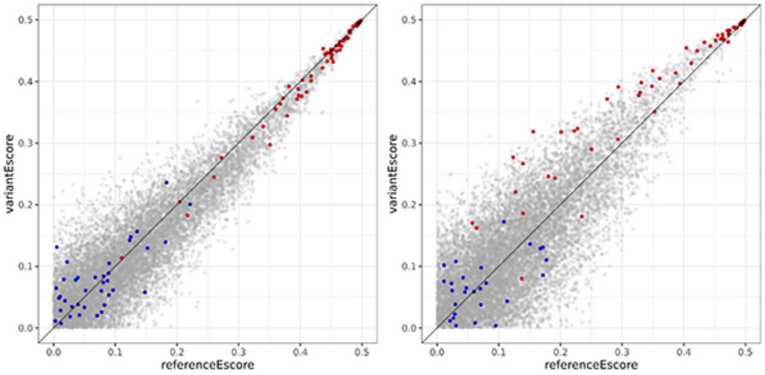


**AB.** FOXF1-Y89C

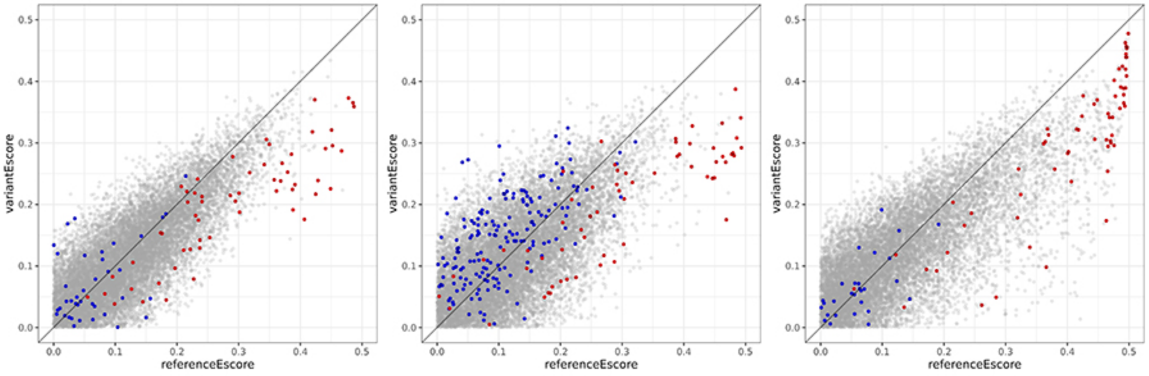


FKH 8-mer
FHL 8-mer
other 8-mer

**Supplementary Figure 1 (cont.)**

**AC.** FOXG1-A188G

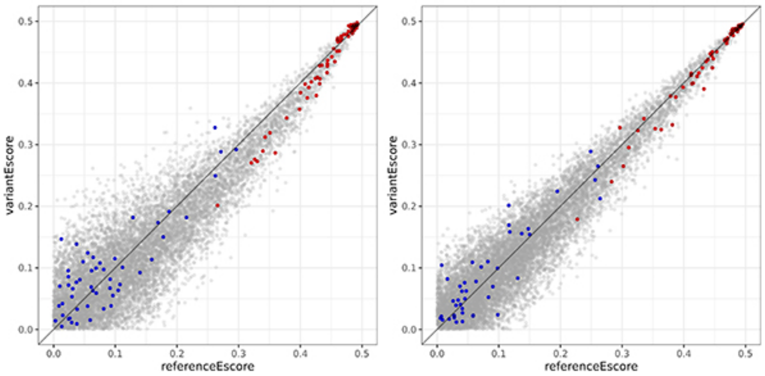


**AD.** FOXG1-R244C

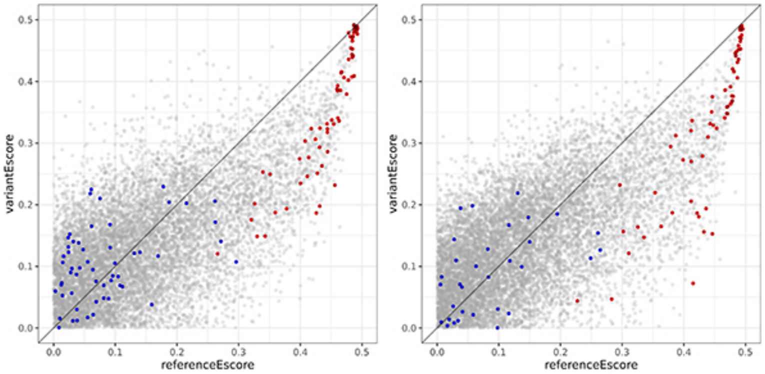


**AE.** FOXG1-R244H

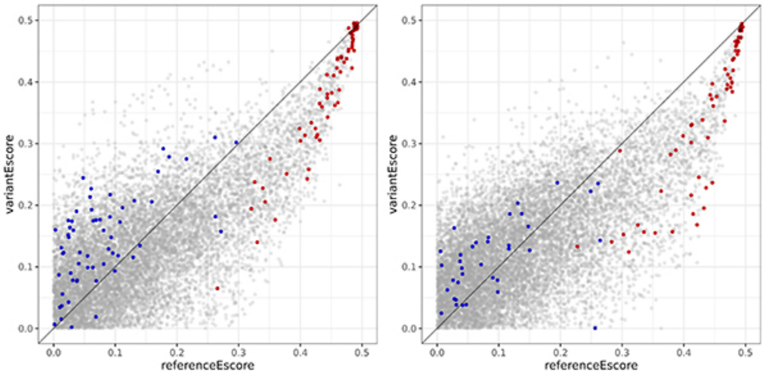


**AF.** FOXG1-R274Q

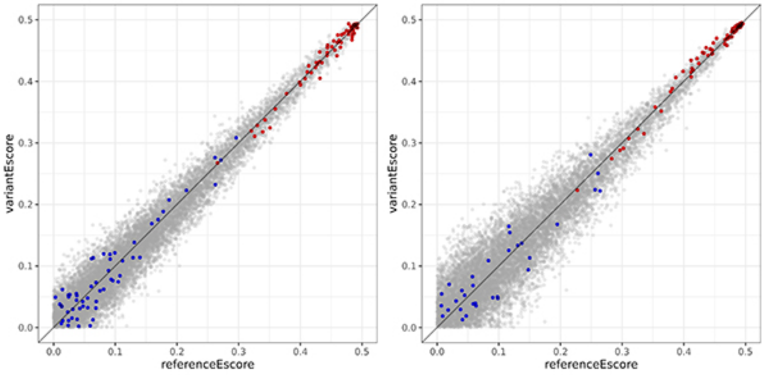


FKH 8-mer
FHL 8-mer
other 8-mer

### Supplementary Figure 1 (cont.)

**AG.** FOXG1-S197I

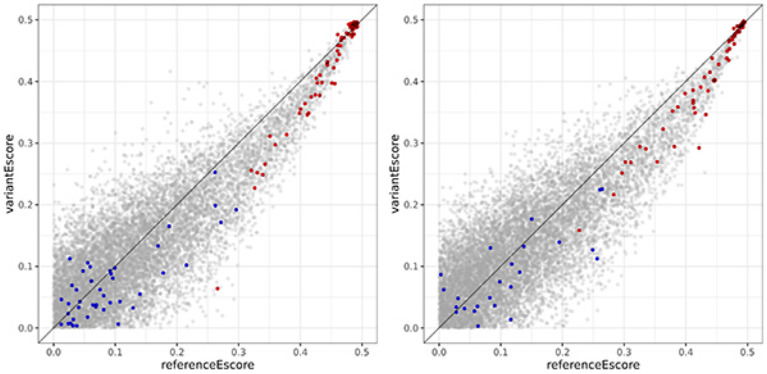


**AH.** FOXI2-D166V

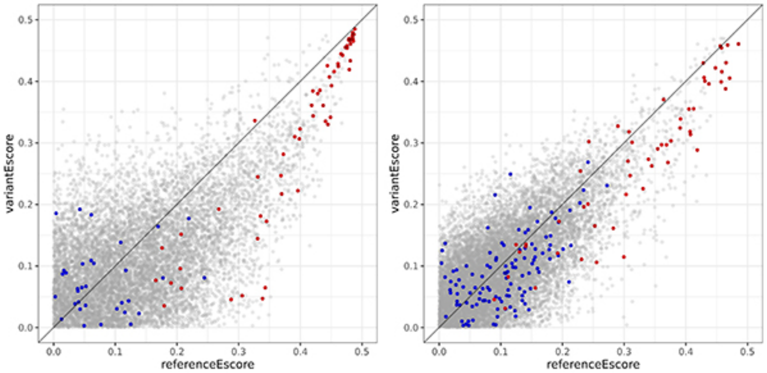


**AI.** FOXI2-D168V

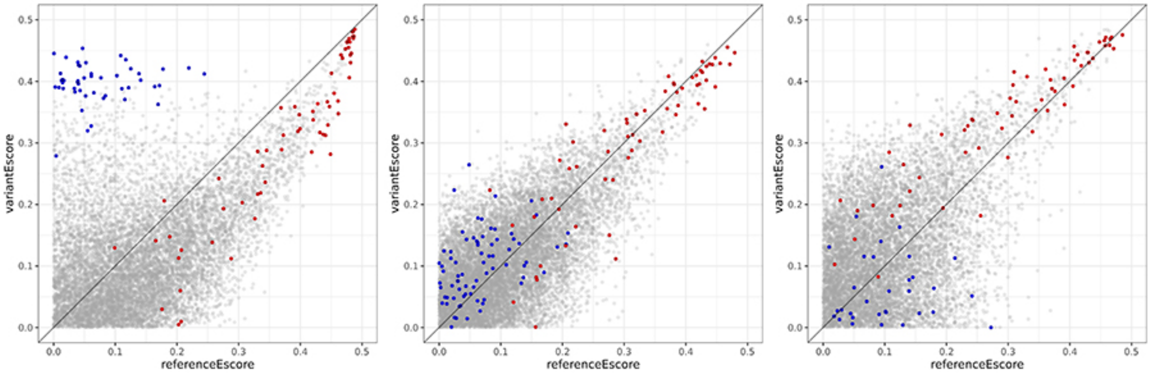


**AJ.** FOXI2-D169E

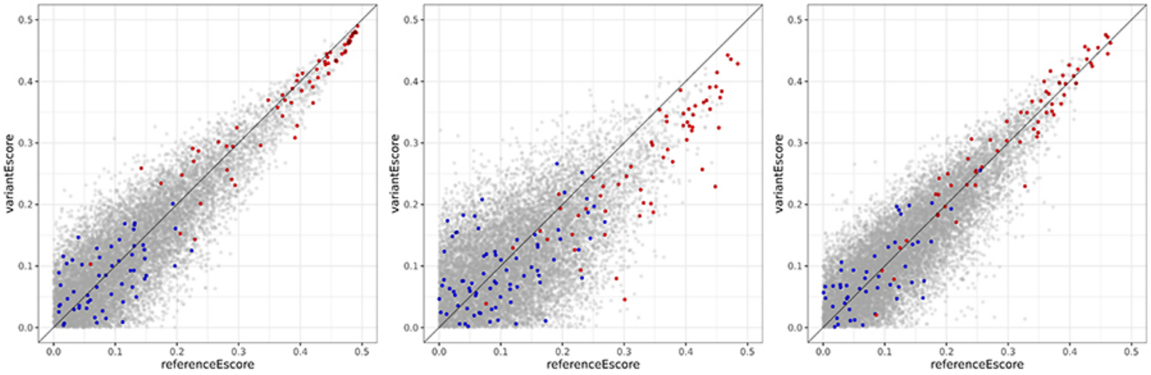


- FKH 8-mer
- FHL 8-mer
- other 8-mer

**Supplementary Figure 1 (cont.)**

**AK.** FOXI2-E200K

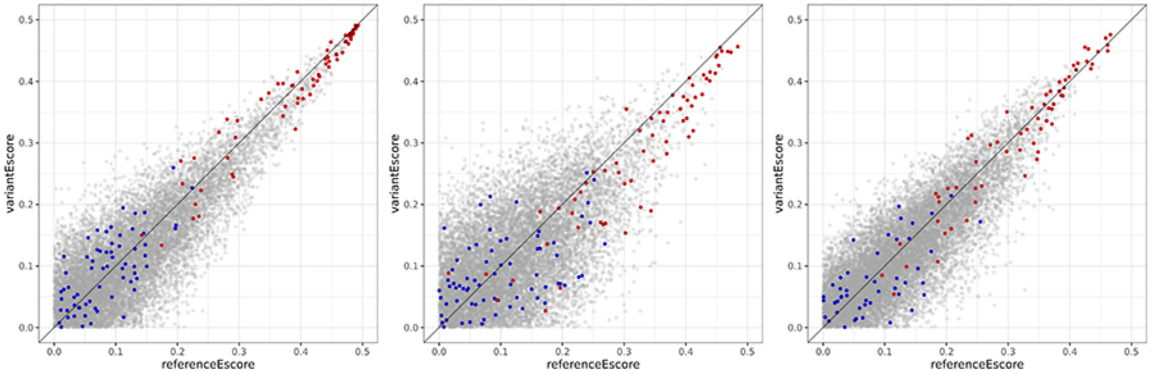


**AL.** FOXI2-G171A

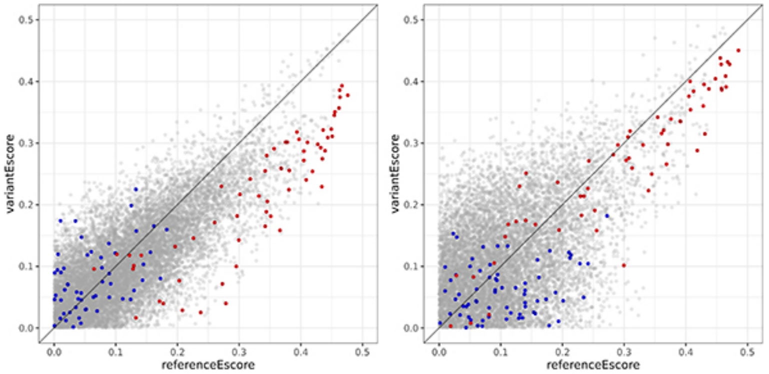


**AM.** FOXI2-G171S

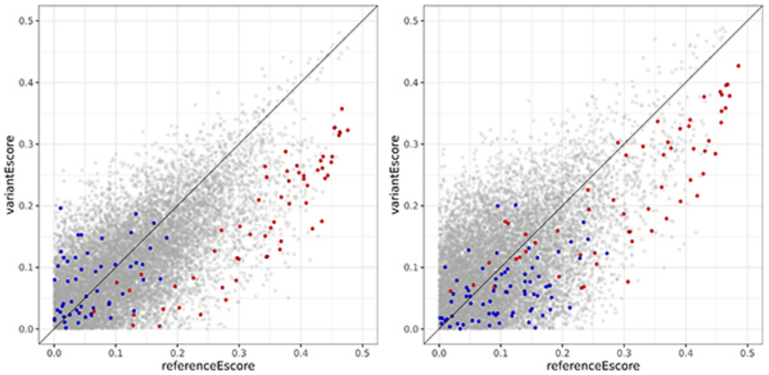


**AN.** FOXI2-K140R

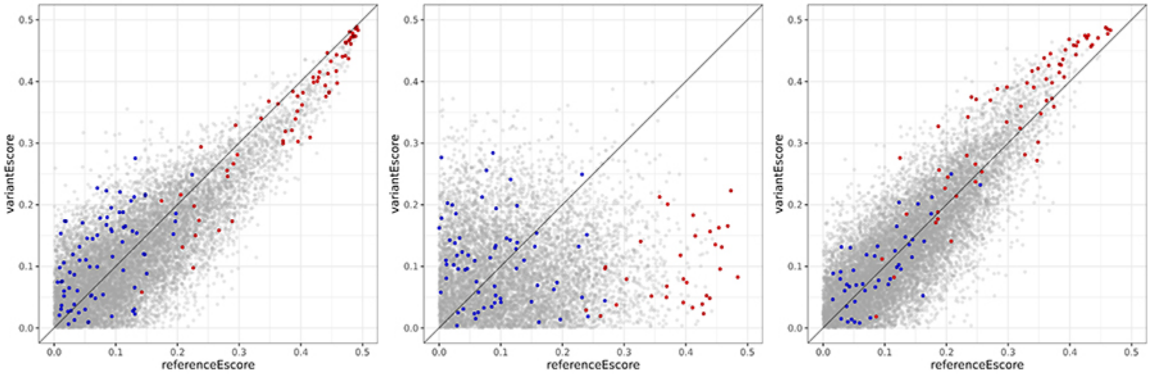


FKH 8-mer
FHL 8-mer
other 8-mer

**Supplementary Figure 1 (cont.)**

**AO.** FOXI2-K184R

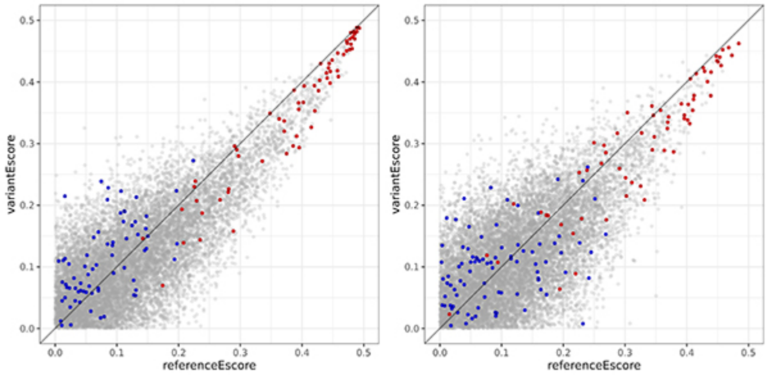


**AP.** FOXI2-L178P

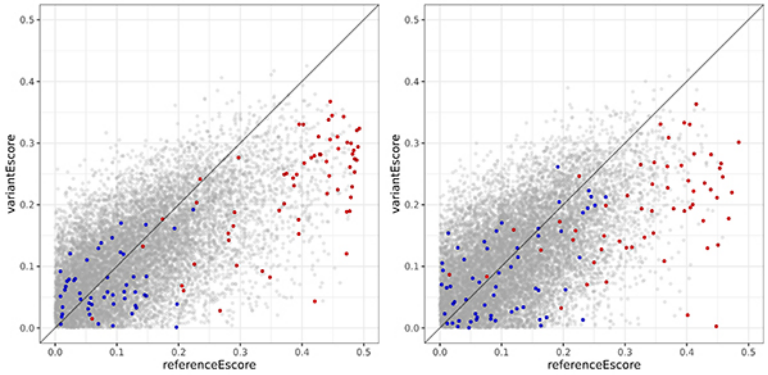


**AQ.** FOXI2-R141L

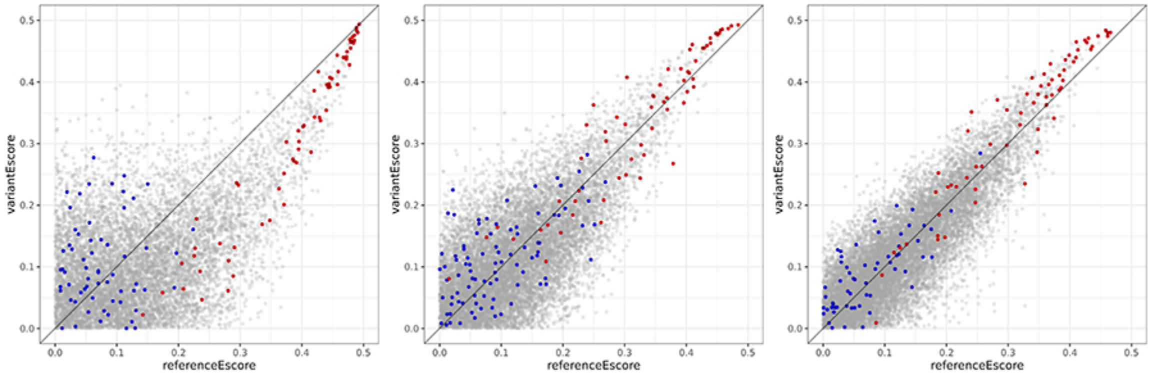


**AR.** FOXI2-W88L

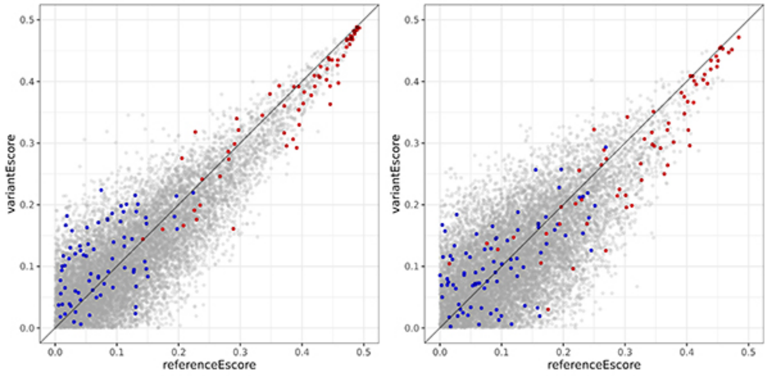


FKH 8-mer
FHL 8-mer
other 8-mer

**Supplementary Figure 1 (cont.)**

**AS.** FOXJ3-H1face

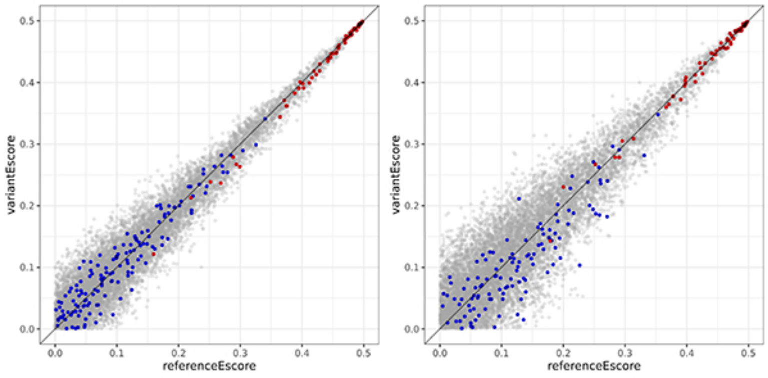


**AT.** FOXJ3-K98N

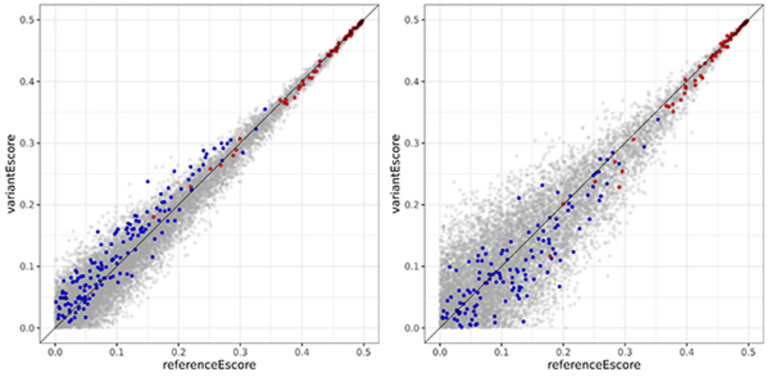


**AU.** FOXJ3-N3wing

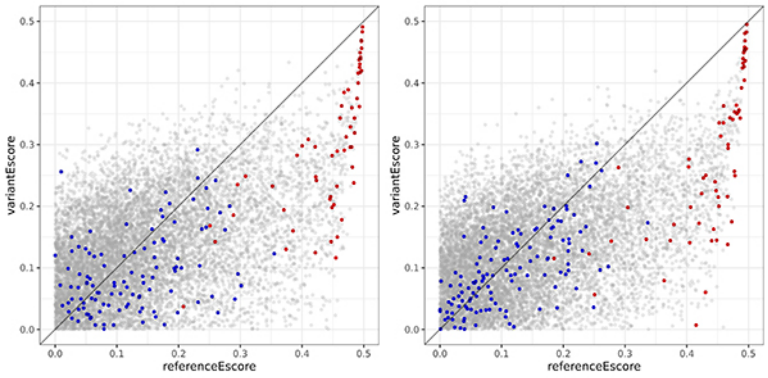


**AV.** FOXJ3-N3wing+H1face

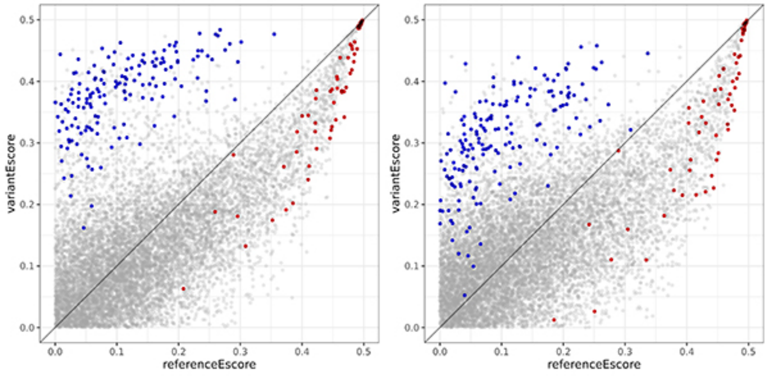


● FKH 8-mer
● FHL 8-mer
● other 8-mer

**Supplementary Figure 1 (cont.)**

**AW.** FOXJ3-V162A

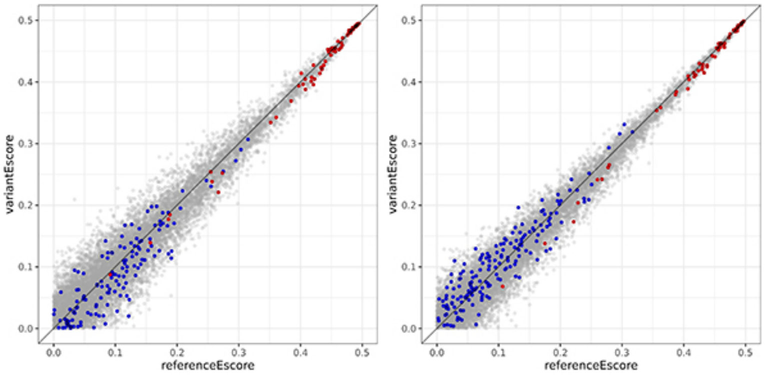


**AX.** FOXN1-D313N

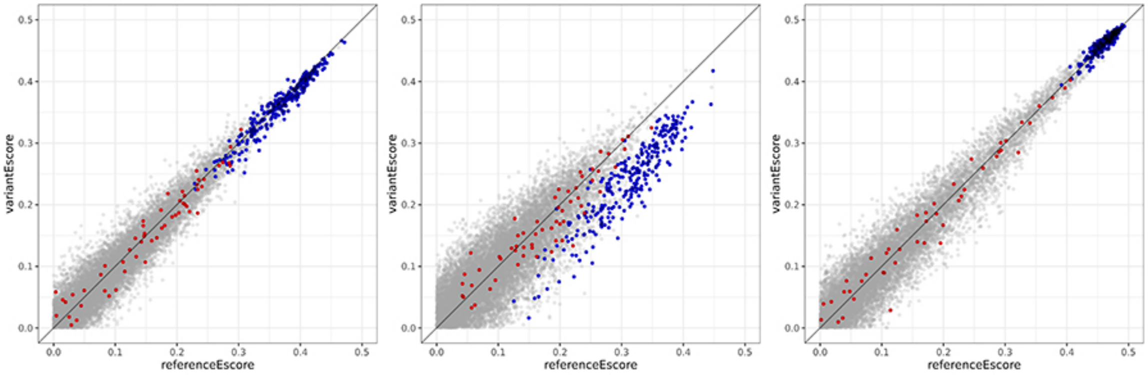


**AY.** FOXN1-E333K

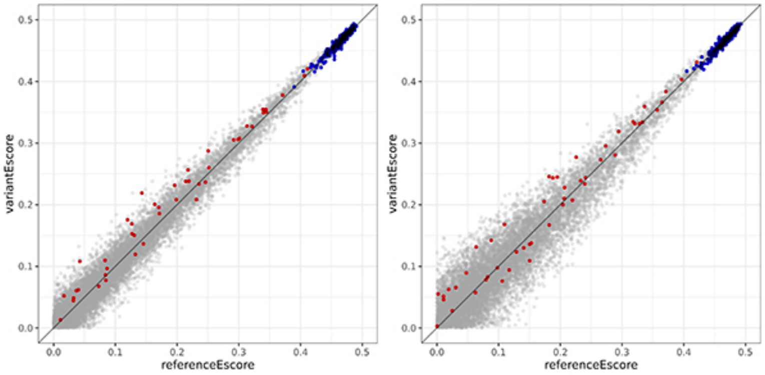


**AZ.** FOXN1-E359K

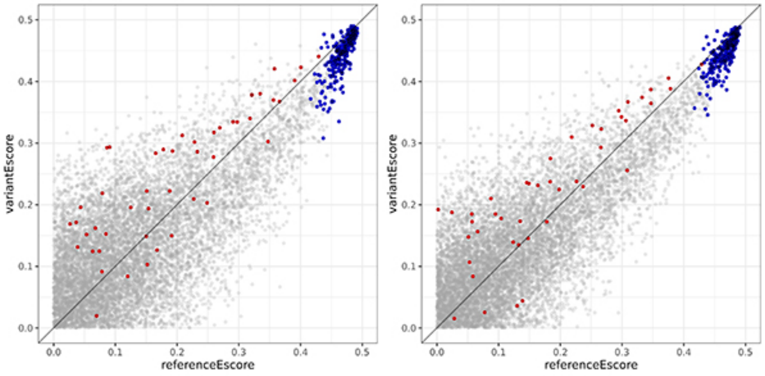


● FKH 8-mer
● FHL 8-mer
● other 8-mer

**Supplementary Figure 1 (cont.)**

**BA.** FOXN1-I278V

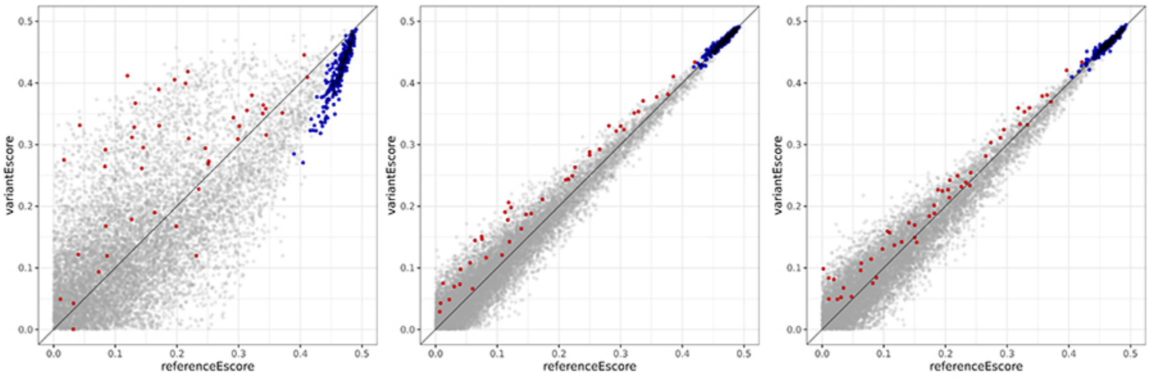


**BB.** FOXN1-N3loop

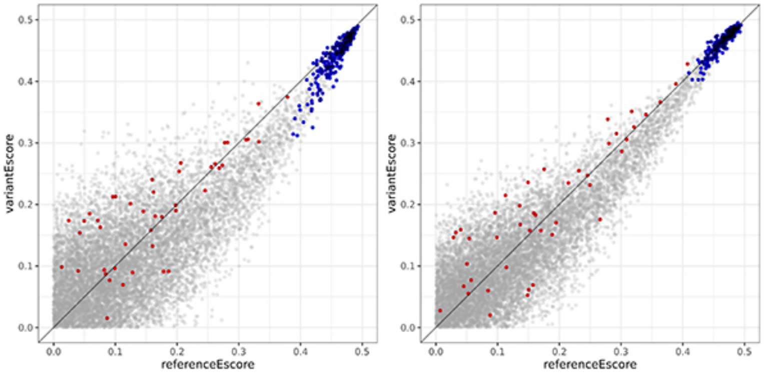


**BC.** FOXN2-K179E

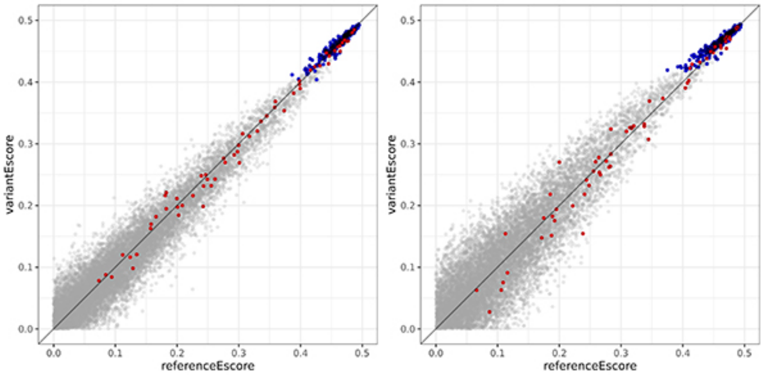


**BD.** FOXN2-L119P

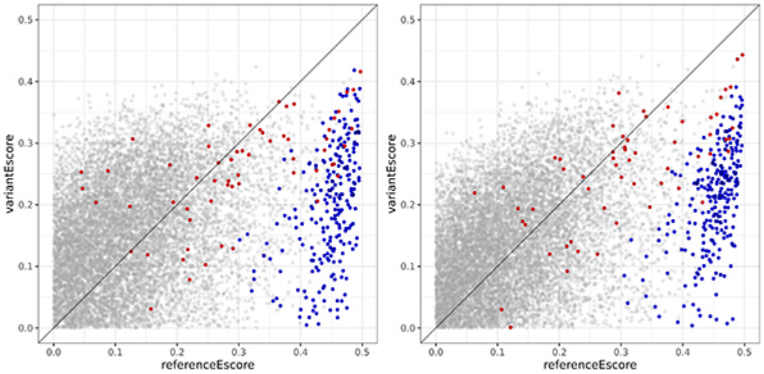


FKH 8-mer
FHL 8-mer
other 8-mer

**Supplementary Figure 1 (cont.)**

**BE.** FOXN2-P153L

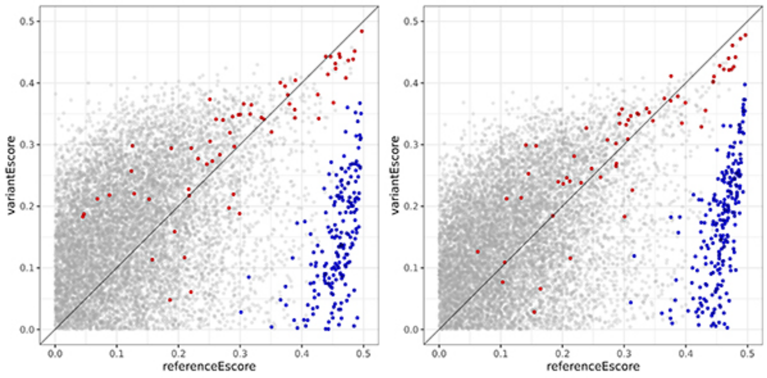


**BF.** FOXN2-T151A

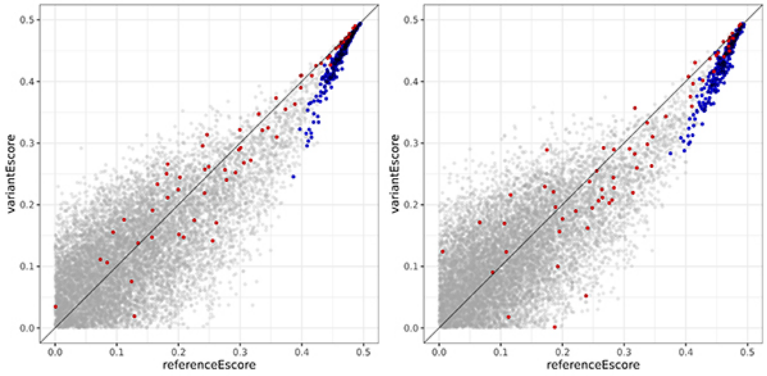


**BG.** FOXN2-T151N

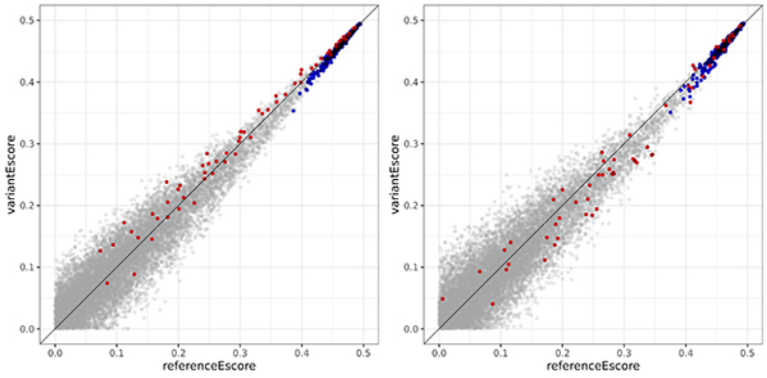


**BH.** FOXN2-T154A

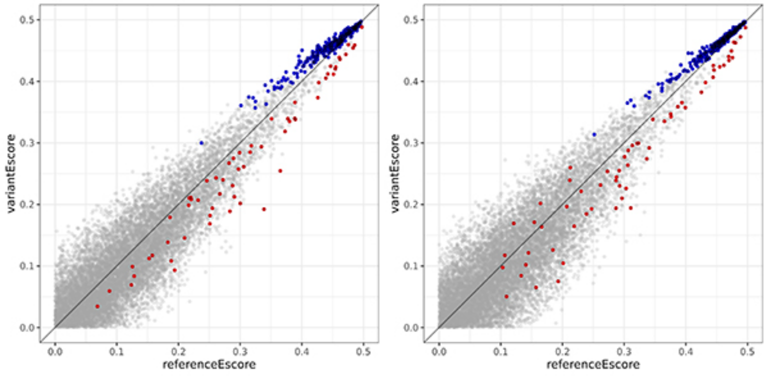


● FKH 8-mer
● FHL 8-mer
● other 8-mer

**Supplementary Figure 1 (cont.)**

**BI.** FOXN3-A1loop

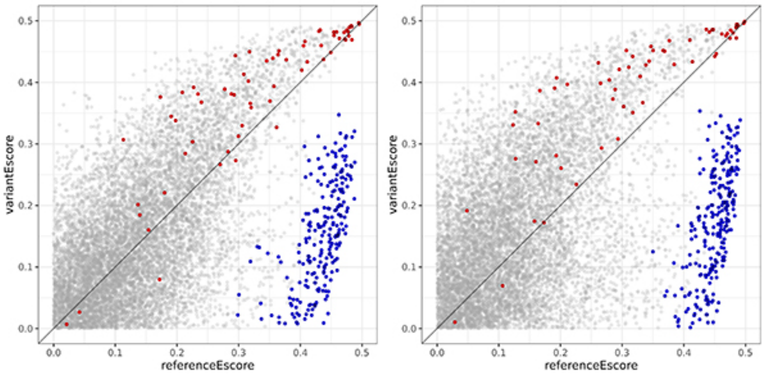


**BJ.** FOXN3-J3wing

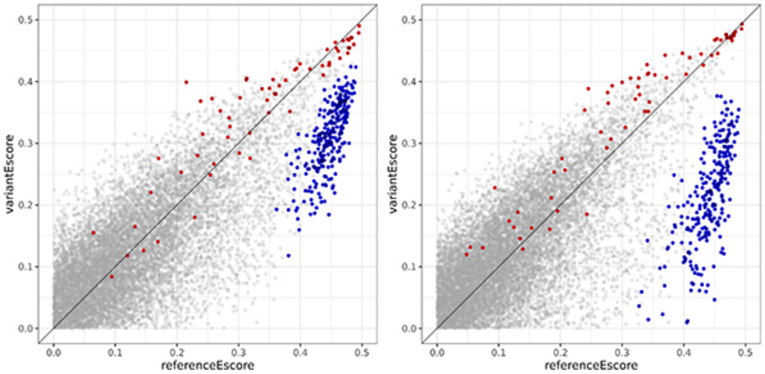


**BK.** FOXN3-J3wing+H1face

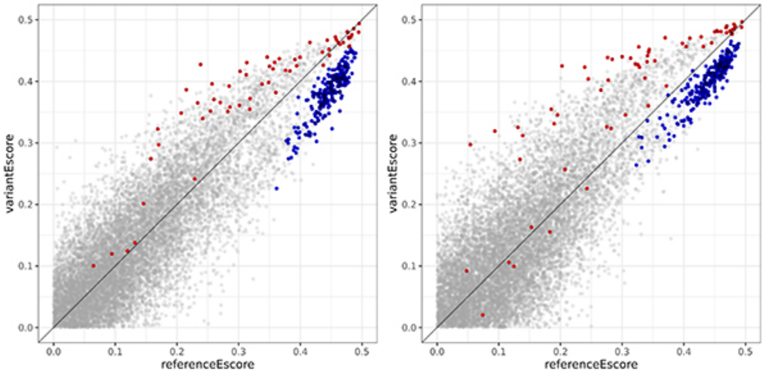


**BL.** FOXN3-L199V

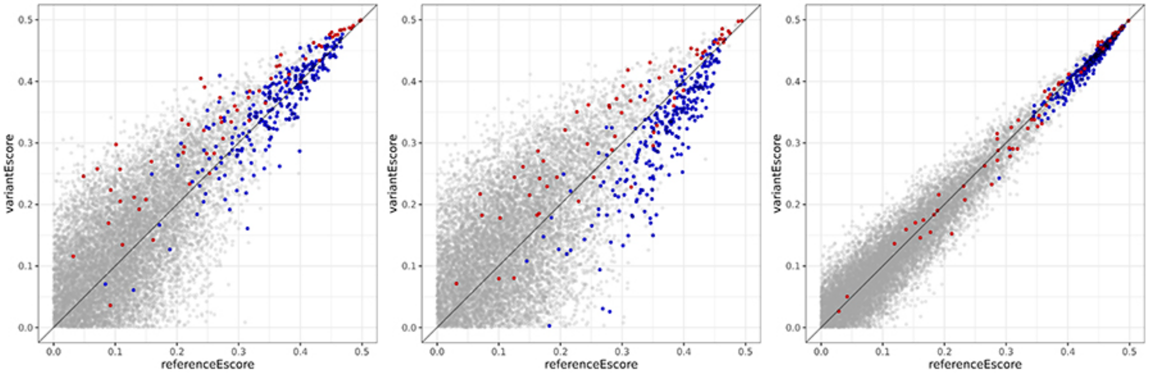


- FKH 8-mer
- FHL 8-mer
- other 8-mer

**Supplementary Figure 1 (cont.)**

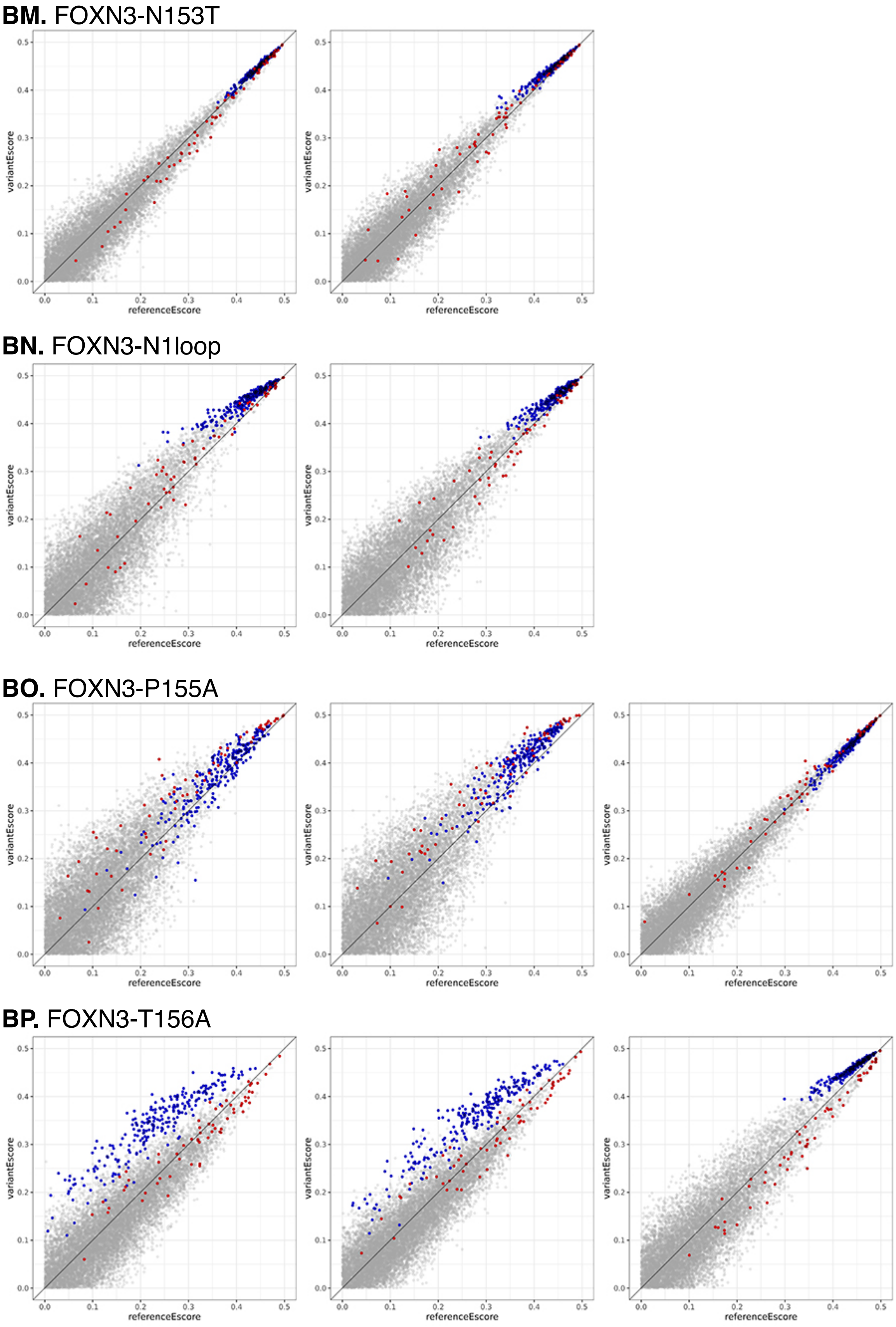


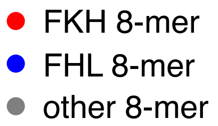


**Supplementary Figure 1 (cont.)**

**BQ.** FOXN3-T156D

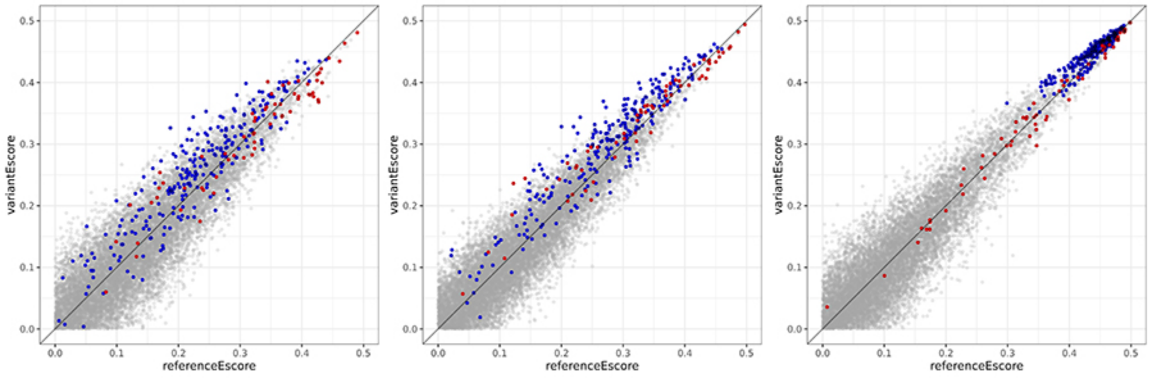


**BR.** FOXN3-T156I

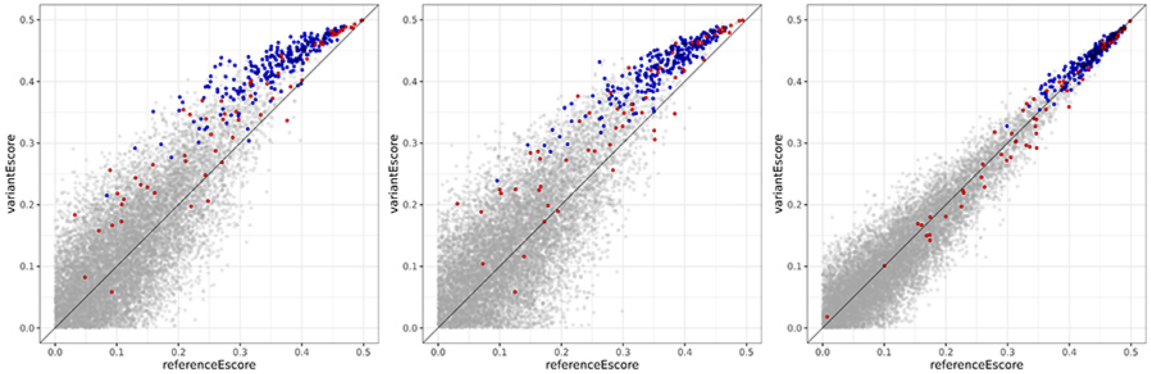


**BS.** FOXN3-T156S

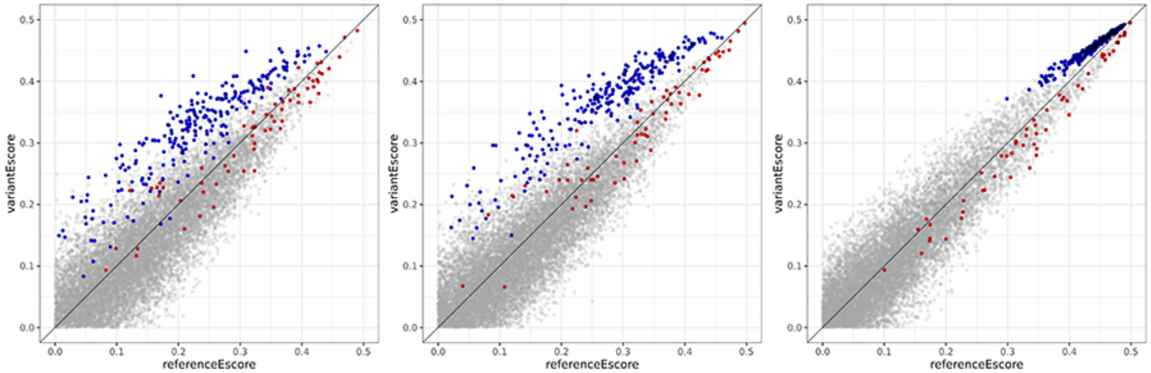


**BT.** FOXN4-G265V

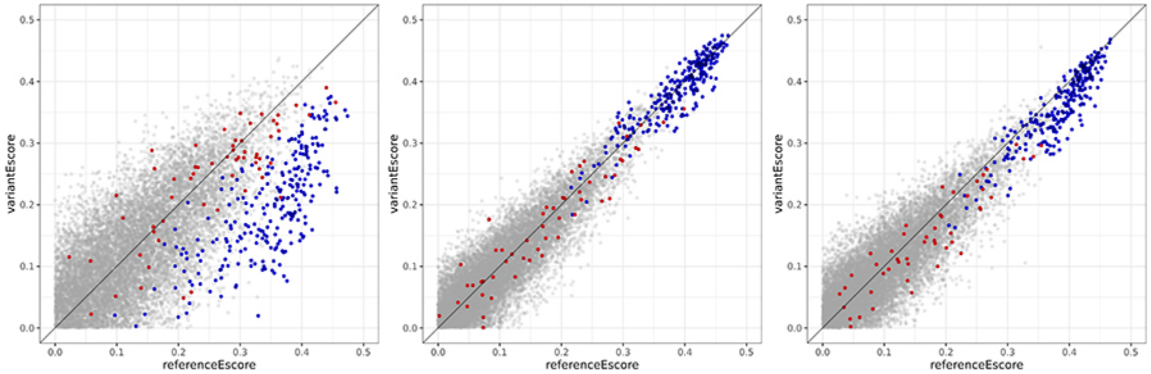


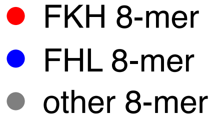


**Supplementary Figure 1 (cont.)**

**BU.** FOXO3-H1face

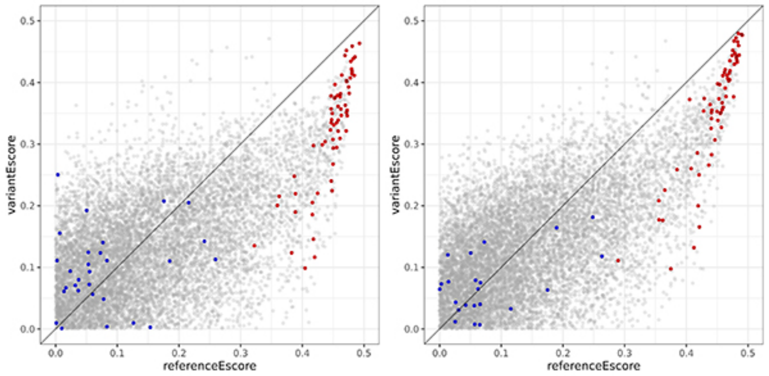


**BV.** FOXO3-N3wing

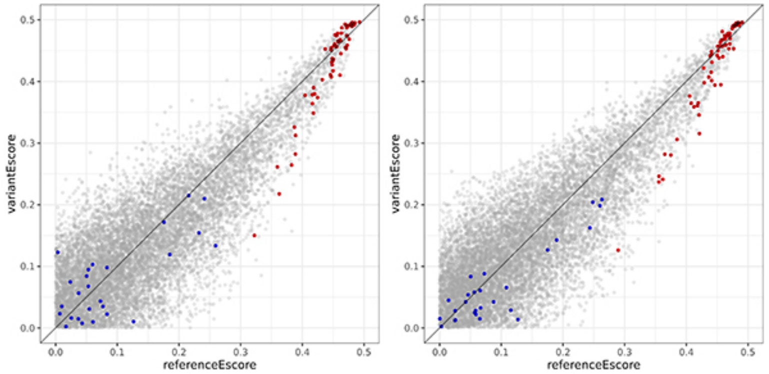


**BW.** FOXO3-N3wing+H1face

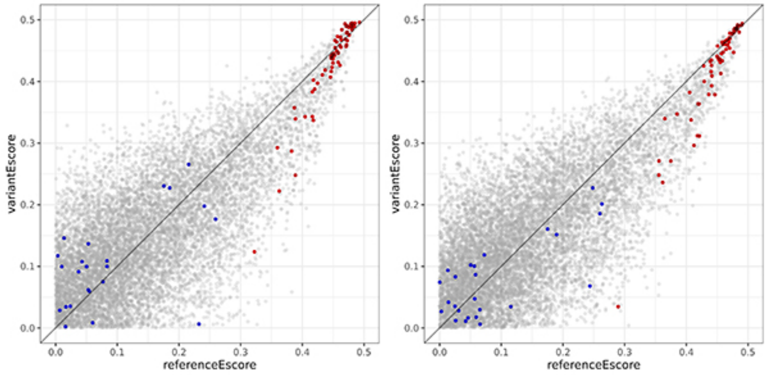


**BX.** FOXS1-H1face

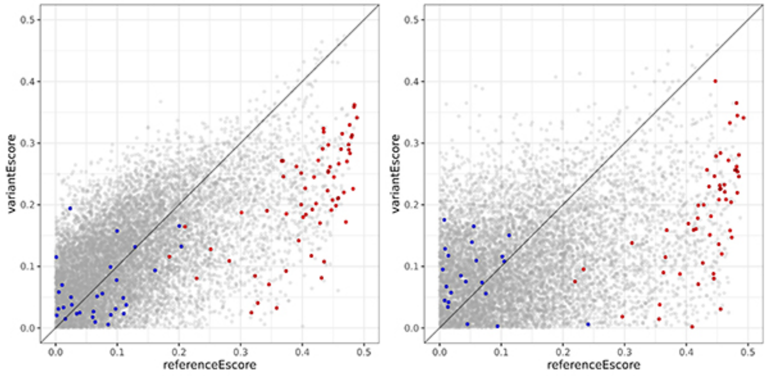


● FKH 8-mer
● FHL 8-mer
● other 8-mer

**Supplementary Figure 1 (cont.)**

**BY.** FOXS1-A28P

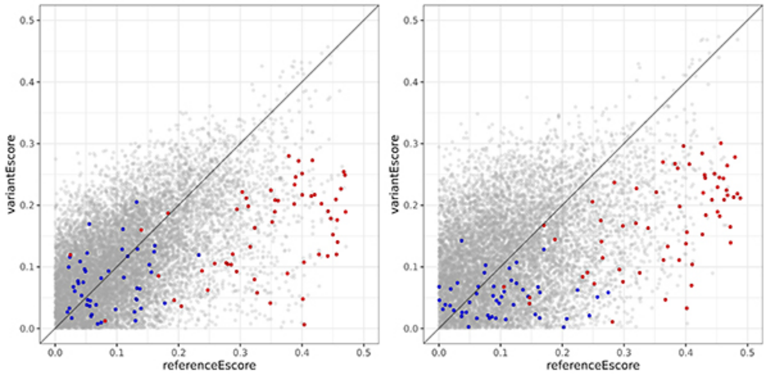


**BZ.** FOXS1-D82G

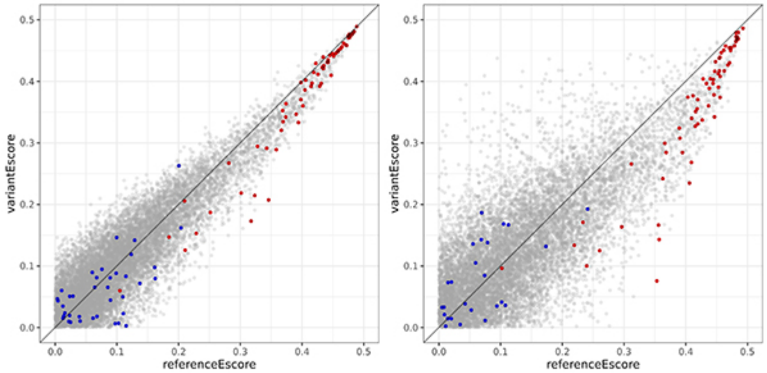


**CA.** FOXS1-K88R

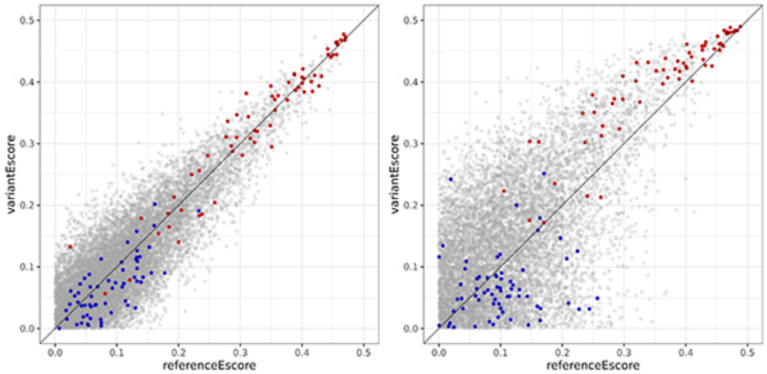


**CB.** FOXS1-N3wing

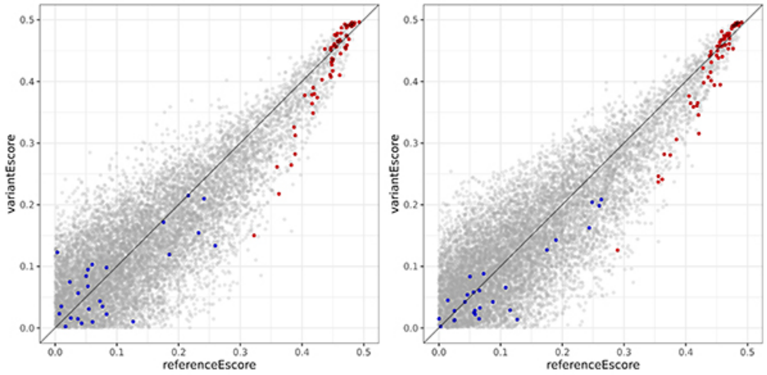


FKH 8-mer
FHL 8-mer
other 8-mer

**Supplementary Figure 1 (cont.)**

**CC.** FOXS1-N3wing+H1face

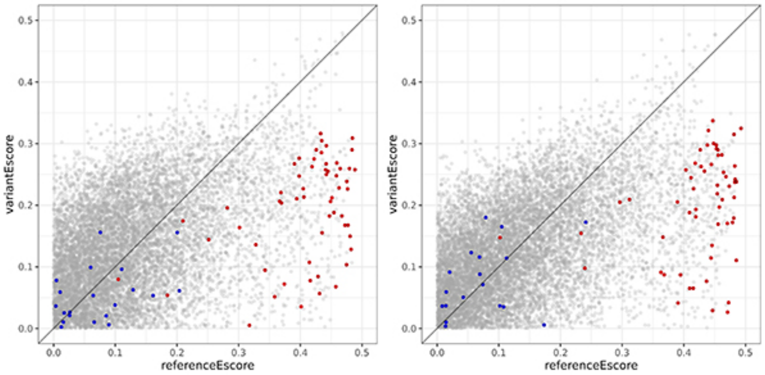


**CD.** FOXS1-R38W

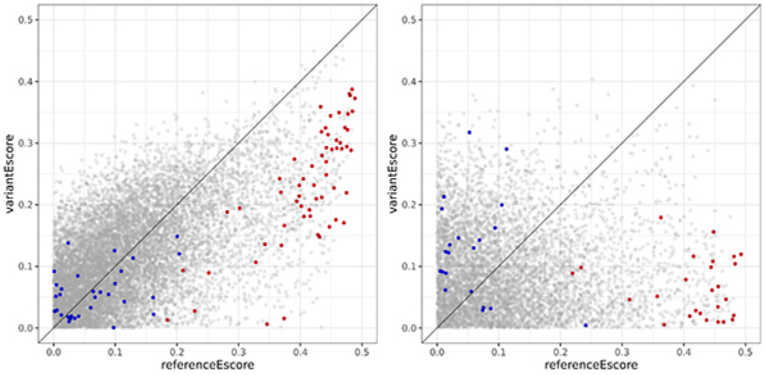


**CE.** FOXS1-R81C

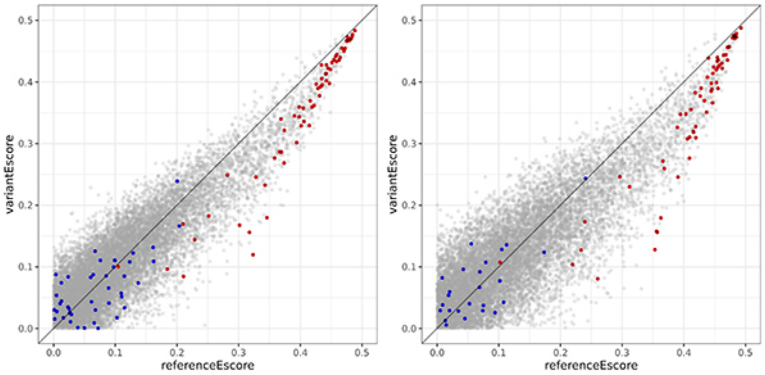


FKH 8-mer
FHL 8-mer
other 8-mer

**Supplementary Figure 1 (cont.)**
